# A plant host enables the heterologous production and combinatorial study of fungal lignin-degrading enzymes

**DOI:** 10.1101/794834

**Authors:** Nikita A. Khlystov, Yasuo Yoshikuni, Samuel Deutsch, Elizabeth S. Sattely

**Affiliations:** Department of Chemical Engineering, Stanford University, Stanford, CA 94305; U.S. Department of Energy Joint Genome Institute, Lawrence Berkeley National Laboratory, Berkeley, CA 94720; Howard Hughes Medical Institute, Stanford University, Stanford, CA 94305

## Abstract

Lignin has significant potential as an abundant and renewable source for commodity chemicals yet remains vastly underutilized. Efforts towards engineering a biochemical route to the valorization of lignin are currently limited by the lack of a suitable heterologous host for the production of lignin-degrading enzymes. Here, we show that expression of fungal genes in *Nicotiana benthamiana* enables production of members from seven major classes of enzymes associated with lignin degradation (23 of 35 tested) in soluble form for direct use in lignin activity assays. We combinatorially characterized a subset of these enzymes in the context of model lignin dimer oxidation, revealing that fine-tuned coupling of peroxide-generators to peroxidases results in more extensive C-C bond cleavage compared to direct addition of peroxide. Comparison of peroxidase isoform activity revealed that the extent of C-C bond cleavage depends on peroxidase identity, suggesting that peroxidases are individually specialized in the context of lignin oxidation. We anticipate the use of *N. benthamiana* as a platform to rapidly produce a diverse array of fungal lignin-degrading enzymes will facilitate a better understanding of their concerted role in nature and unlock their potential for lignin valorization, including within the plant host itself.

## Main

The utilization of lignin as a highly abundant renewable feedstock for platform commodity chemicals remains a long-sought goal for over four decades^1^. 1.3 billion tons of lignocellulosic biomass are available each year in the U.S. alone, of which lignin comprises up to 40% by weight^1^. The lack of an established route to lignin deconstruction means that the primary value of industrial lignin by-products is heat and electricity by incineration. Molecular decomposition of lignin would yield monomeric aromatic derivatives with economic value in chemical and material applications typically reliant on petroleum, with vanillin^2^ and phenolic resins^3^ as prominent examples.

Biological catabolism of lignin has been widespread in nature for at least 200 million years^4^, presenting scalable means of its deconstruction through renewable enzymatic catalysis under mild conditions. Wood-decaying fungi are particularly proficient at lignin catabolism and are regarded as the primary agents of lignin biodegradation in nature^5^. Some, such as the model white-rot basidiomycete *Phanerochaete chrysosporium*, have been reported to be capable of degrading 1 *g* lignin per *g* mycelium per day^6^. Genomic sequencing of this model species as well as other wood-decaying relatives has uncovered a diverse assortment of genes involved in lignin decomposition^7,8^, making basidiomycetes attractive potential sources of wood-decaying enzymes for biotechnological applications. However, basidiomycetes remain genetically intractable and challenging to cultivate rapidly, rendering their utility in the industrial production of lignin-degrading enzymes unrealized.

Efficient heterologous production of basidiomycete lignin-degrading enzymes has also proven elusive^9,10^. Lignin biodegradation in nature is largely driven by the oxidative action of heme peroxidases secreted by white-rot basidiomycetes^4,11^. Heme cofactor incorporation, disulfide bond formation, and glycosylation have rendered this class of enzymes particularly challenging to over-express heterologously^10,12,13^. Model bacterial expression platforms such as *Escherichia coli* are poorly suited for the required post-translational features of these enzymes^14^. Previous characterization of this class of enzymes has relied primarily on *in vitro* refolding from bacterial inclusion bodies, an inherently time-consuming and inefficient process^9^. While yeasts such as *Saccharomyces cerevisiae* present a logical choice of model eukaryotic microbial hosts, they do not natively secrete heme peroxidases. Therefore, the development of an expression host comprehensively capable of heterologous production of lignin-degrading enzymes is of paramount importance in establishing a biochemical route to lignin valorization^9^.

To this end, we hypothesized that a plant-based expression system could serve as an effective chassis for the heterologous production of basidiomycetes lignin-degrading enzymes. Plants produce numerous extracellular heme peroxidases for cell wall biosynthesis and morphogenesis, with the widely used experimental host *Nicotiana benthamiana*, a close relative of tobacco, natively producing six class III peroxidases^15,16^. Because of its amenability to protein over-expression through well-established *Agrobacterium* infiltration methods for DNA delivery^17^ and its native capacity to secrete heme peroxidases, we reasoned that *N. benthamiana* would be well-suited as a heterologous host for the production of fungal lignin-degrading peroxidases.

Basidiomycetes employ a diverse arsenal of enzymes to achieve lignin degradation (**Fig. 1a**), featuring numerous isoforms of individual enzyme types^8^. Given their uniquely high redox potential sufficient to perform one-electron oxidation of the aromatic lignin backbone^18^, we prioritized the study of a large array of members from across the three major white-rot peroxidase families. Lignin peroxidases (LiP) are known to oxidize lignin-related compounds at the enzyme surface through long-range electron transfer (LRET), manganese peroxidases (MnP) through oxidation of Mn(III) as a diffusible mediator, and versatile peroxidases (VP) through both modes of oxidation^19^. As these peroxidases use peroxide as a co-substrate, we also selected a number of basidiomycete peroxide-generating enzymes from the pyranose oxidase (POx) and aryl alcohol oxidase (AAO) families to characterize coupled activity alongside the peroxidases. To explore the general capacity of heterologous hosts to produce lignin-degrading enzymes, we also selected members of the laccase (Lac) and cellobiose dehydrogenase (CDH) enzyme families, which are thought to catalyze the oxidation of phenolic lignin fragments and to interface with cellulose deconstruction, respectively^20^.

**Figure 1.**
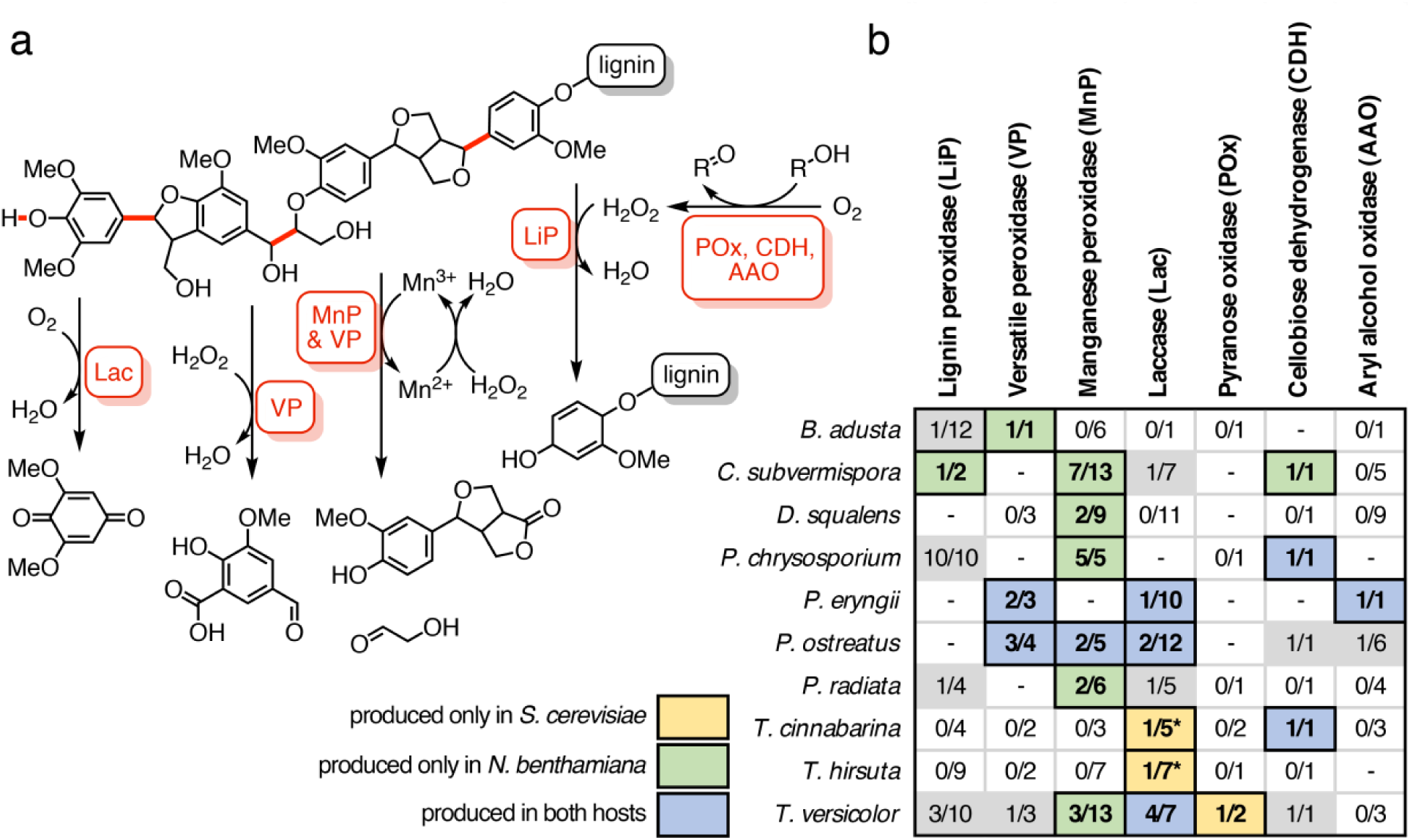
**a)** A representative schematic of bond cleavage catalyzed by different families of ligninases. Featured in red are carbon-carbon bonds that can be cleaved by different types of heme peroxidases (MnP, VP, LiP), as well as phenolic bonds that can be cleaved by laccases (Lac). Auxiliary enzymes such as aryl alcohol oxidase (AAO), cellobiose dehydrogenase (CDH), and pyranose oxidase (POX) are shown as examples that can be coupled to peroxidase activity on lignin bonds. **b)** An overview of ligninases tested in this study. Each entry indicates the number of isozymes tested out of the total number of isozymes native to each fungal species. Entries bolded and highlighted in yellow indicate successful heterologous production in *S. cerevisiae*, those in green for *N. benthamiana*, and those in blue for both species of one or more isozymes in that category. Entries with dashes indicate absence of that enzyme type in the fungal species, and those having zero as the first number indicate that no representative genes from that species and enzyme class were synthesized or tested. Laccases from *T. cinnabarina* and *T. hirsuta* were tested only in *S. cerevisiae* and are denoted with an asterisk.

While individual members of the peroxidase families have been shown to catalyze a reduction in the molecular weight of synthetic lignin *in vitro*^21,22^, many open questions remain surrounding lignin deconstruction in nature. Why do white-rot basidiomycetes possess as many as 26 genes encoding peroxidases^8^, including 13 manganese peroxidase isoforms^23^, and how are their activities coordinated during lignin degradation? To begin to answer these questions, we tested the production of an extensive panel of members from the major classes of lignin-degrading enzymes in both the model eukaryotic microbial host *S. cerevisiae* and the model plant *N. benthamiana*. Our results suggest that *N. benthamiana* is better suited for the production of lignin-degrading enzymes and fulfils the role of an effective production platform required for their study, with 23 of 35 enzymes tested expressed in active form. Combinatorial assay of a selected set of successfully produced enzymes for cleavage of a model lignin dimer revealed that the extent of lignin C-C bond scission is enhanced in a coupled system, as well as by the action of specific isozymes compared to others from a given fungal species. Taken together, these results provide insight into the specialization of different isozymes and will help facilitate an effective means of engineering biochemical lignin valorization.

## Results

### *Improved heterologous production of lignin-degrading enzymes* in plantae

Our proposed approach to examine the potential synergistic activity of lignin-degrading enzymes (**Fig. 1a**) requires a comprehensive set of basidiomycete proteins for combinatorial analysis (**Fig. 1b**). Given the genetic intractability and the native lignolytic background of basidiomycetes, we opted for heterologous production to obtain these enzymes, attempting to do so in *S. cerevisiae* first. Production of enzymes in soluble form without refolding was desirable for rapid screening and required a eukaryotic host to achieve proper folding and post-translational modification of the secreted lignin-degrading enzymes. Sourcing from annotated fungal genomes, we identified 65 genes with previous transcriptomic, secretomic and/or biochemical characterization spanning seven major classes of lignin-degrading enzymes. We codon-optimized and *de novo* synthesized genes encoding for mature enzymes fused to a variety of yeast-based signal peptides and affinity tags, cloning into low- and high-copy expression cassettes (**Fig. 2a**). We observed that a strain background repaired of mitochondrial defects^24^ was more proficient in secretion of a model peroxidase compared to the commonly-used secretion strain BJ5465^25^ (**Fig. S1**) and used this strain for heterologous production of our array of fungal genes. We also tested an array of genetic components, media supplements, and growth temperatures (**Figs. S1 and S2**) and arrived at a high-copy expression cassette (**Fig. 2a**) and cultivation conditions best-suited for high-throughput screening of enzyme production in *S. cerevisiae*.

**Figure 2.**
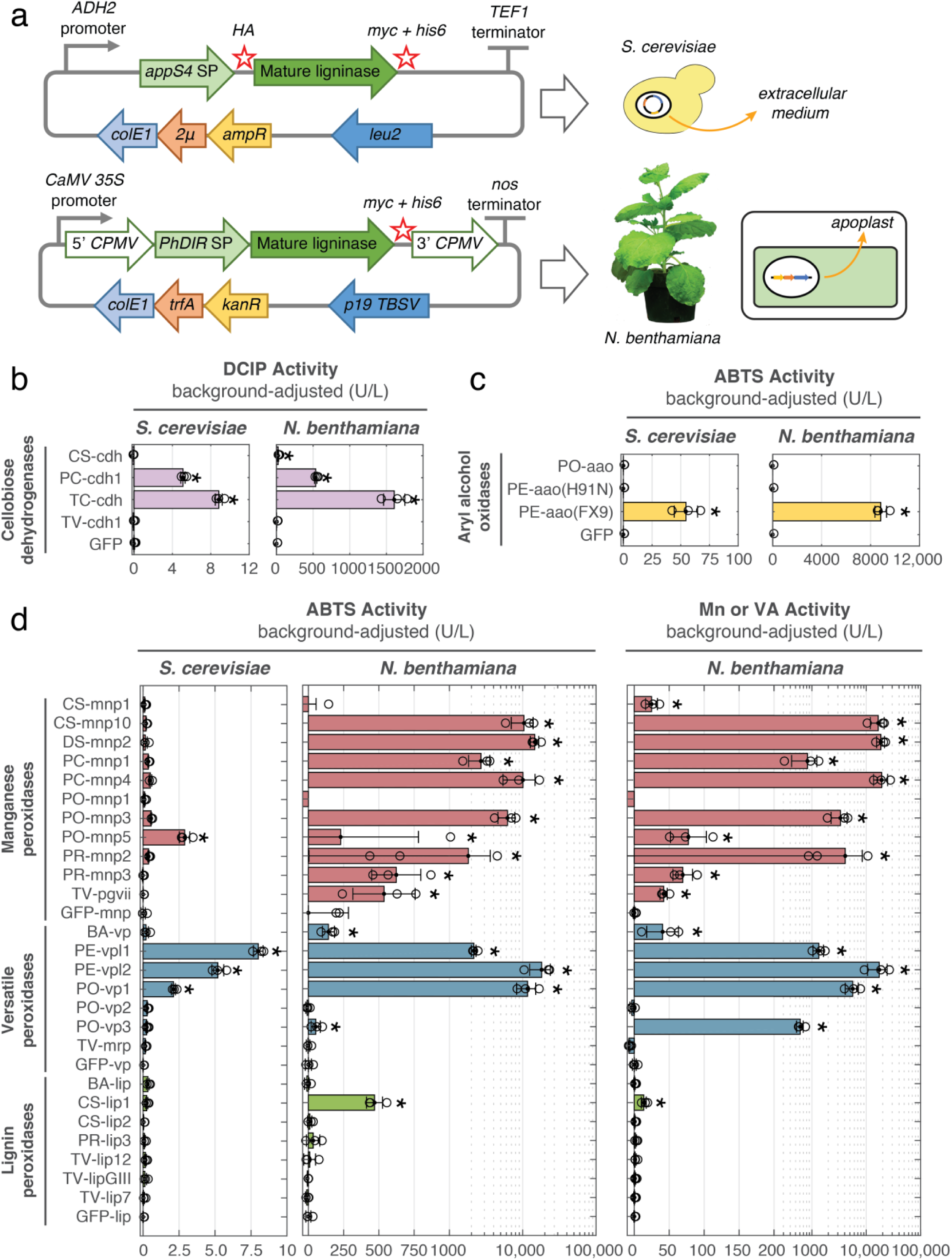
**a)** Overview of heterologous production of lignin-degrading enzymes in *S. cerevisiae* and *N. benthamiana*. For *S. cerevisiae*, production of enzymes was driven by an ADH2 promoter on a pCHINT2AL high-copy expression vector^24^ and exported to the extracellular medium (supernatant) via fusion to an evolved variant of *S. cerevisiae* α-mating-factor signal peptide (*appS4 SP*)^28^. Production of enzymes in *N. benthamiana* was driven by a 35S promoter on a pEAQ expression vector^29^ and exported to the plant apoplast via fusion to the signal peptide of dirigent protein from *Sinopodophyllum hexandrum* (PhDIR SP). The pEAQ vector includes 5’ and 3’ untranslated regions from cowpea mosaic virus (*5’ CPMV and 3’ CPMV*) to enhance expression levels. Hemaaglutinin (*HA*), c-myc (*myc*), and hexahistidine (*his6*) affinity tags were included as N- and/or C-terminal fusions where indicated. **b)** Activity of cellobiose dehydrogenases expressed in *S. cerevisiae* and *N. benthamiana*, measured spectroscopically as DCIP reduction. **c)** Activity of aryl alcohol oxidases expressed in *S. cerevisiae* and *N. benthamiana*, measured spectroscopically as HRP-coupled oxidation of ABTS in the presence of veratryl alcohol (VA). **d)** Summary of lignin-degrading heme peroxidase activity in culture supernatant of *S. cerevisiae* or in apoplast extracts of *N. benthamiana*, assayed against ABTS and VA^30^ or Mn(II)^31^. Lignin peroxidases are shown in green, versatile peroxidases in blue, and manganese peroxidases in red. Mn/VA activity corresponds to Mn activity (1 mM MnSO_4_, 100 µM H_2_O_2_) for manganese and versatile peroxidases and VA activity (20 mM veratryl alcohol, 100 µM H_2_O_2_) for lignin peroxidases. ABTS activity assays (4 mM ABTS, 100 µM H_2_O_2_) of manganese peroxidase samples also included 1 mM MnSO_4_. Activities represent 1 µM oxidized product formed *min*_-1_ *l*_-1_. Activities were measured of three different leaves in *N. benthamiana* 5 days after *Agrobacterium* infiltration, or three biological replicates for *S. cerevisiae* after 2 days of cultivation. Reported activities are measured activity levels minus background activity levels in the corresponding GFP sample for each set of enzymes. Asterisks indicate statistically significant activity levels relative to GFP control samples (*p* < 0.05, excluding outliers). BA = *Bjerkandera adusta*, CS = *Ceriporiopsis (Gelatoporia) subvermispora*, PC = *Phanerochaete chrysosporium*, PE = *Pleurotus eryngii*, PO = *Pleurotus ostreatus*, PR = *Phlebia radiata*, TC = *Trametes (Pycnoporus) cinnabarinus*, TV = *Trametes versicolor*.

Using this optimized expression platform, we successfully produced six of 11 laccases, a pyranose oxidase, and two of four cellobiose dehydrogenases using *S. cerevisiae* as indicated by activity assays (**Figs. 2b and S3**). Aryl alcohol oxidase production was achieved through use of a previously-evolved, well-expressing enzyme variant^26^ (**Fig. 2c**). Of greatest interest were the heme peroxidases, since this class of enzymes are considered to be the primary agents of lignin degradation. Out of 47 isozymes, only four were produced to any extent measurable by activity assays towards the model peroxidase substrate 2,2’-azino-bis(3-ethylbenzothiazoline-6-sulphonic acid) (ABTS) (**Fig. 2d and S4**), with no detectable activity towards the lignin-related substrates Mn(II) and veratryl alcohol (*data not shown*). We also observed that yeast media supernatant inhibited the activity of commercially available lignin peroxidase towards veratryl alcohol, suggesting interference of secreted yeast metabolites in oxidation of more recalcitrant substrates by lignin-degrading peroxidases (**Fig. S5**).

Given our interest in testing the activity of lignin-degrading peroxidases, we next attempted their production in *N. benthamiana*, which we hypothesized would be more amenable as a heterologous host given its native repertoire of secreted plant peroxidases. In order to test heterologous production of lignin-degrading enzymes in a high-throughput manner, we employed *Agrobacterium*-mediated transient expression of fungal genes in leaf tissue of mature plants. We targeted the proteins to the plant apoplast via C-terminal fusion to the signal peptide of a dirigent protein from *Sinopodophyllum hexandrum* (**Fig. 2a**) to provide access to post-translational modifications and an appropriate folding environment. This targeting strategy also enabled straightforward extraction of heterologous enzymes in soluble form by centrifugation of leaf tissue^27^ (**Fig. S6**). In contrast to yeast, our approach of using a plant-based host provided a diverse set of isozymes from each of the three major types of white-rot lignin-degrading peroxidases, including 10 manganese peroxidases (MnPs), five versatile peroxidases (VPs), and a lignin peroxidase (LiP) (**Fig. 2d**). Raw apoplast extracts of plants expressing fungal peroxidases displayed high levels of activity towards veratryl alcohol and Mn(II) substrates without inhibition, in contrast to production in *S. cerevisiae* (**Fig. 2d**). Further improvements in activity levels were possible through the use of signal peptides native to *N. benthamiana* (**Fig. S7**). *N. benthamiana* also proved proficient in the production of cellobiose dehydrogenases and an aryl alcohol oxidase (**Figs. 2b and 2c**). In the case of pyranose oxidase, however, no measurable activity was detected in apoplast extracts from *N. benthamiana*, and *S. cerevisiae* was subsequently used for production of this enzyme (**Fig. S3**). In general, *N. benthamiana* greatly exceeded *S. cerevisiae* in activity of enzyme produced (as determined by comparing activity *l*_-1_ plant apoplast extract vs. *l*_-1_ yeast supernatant), and enabled streamlined production of a variety of lignin-degrading enzymes for *in vitro* lignin activity testing.

### Model lignin dimer oxidation

Equipped with a diverse set of heterologously-produced fungal lignin-degrading enzymes, we sought to uncover their coordinated roles in the context of lignin degradation. We selected a β- O-4 dimer as a model representative of the most common type of linkage found in lignin^32^, circumventing the problematic insolubility of whole lignin and enabling rapid product characterization by liquid-chromatography mass spectroscopy (LC-MS) while serving as a robust indicator of the lignin bond cleavage ability of a given enzyme. Given the sensitivity of lignin-degrading peroxidases to suicide inactivation by excess peroxide^33^, we sought to better approximate biological delignification and provide peroxide via an enzyme coupling approach rather than by direct addition of peroxide to help mitigate this issue. This was achieved by coupling peroxidases to peroxide-generating enzymes such as sugar oxidases and aryl alcohol oxidases, thought to be the primary sources of peroxide in wood-decaying basidiomycete fungi^34–37^. We selected a representative from each type of lignin-degrading peroxidase: CS-*lip1* as a lignin peroxidase; PC-*mnp1* as a manganese peroxidase; and PE-*vpl2* as a versatile peroxidase. We chose a pyranose oxidase from *T. versicolor* and produced in *S. cerevisiae*, the evolved variant of aryl alcohol oxidase^26^ from *P. eryngii* and produced in *N. benthamiana*, and commercially-available glucose oxidase from *A. niger* as peroxide-supplying counterparts to the peroxidases in the coupled reactions.

Through kinetic sampling of the reactions by LC-MS, we identified three major products resulting from the oxidation of the model dimer^32^: veratraldehyde and a Hibbert ketone, resulting from carbon-carbon bond cleavage; and dehydrodimer, resulting from oxidation of the α-hydroxy group without bond cleavage (**Fig. 3a**). We observed accumulation of these products in all coupled reactions containing diafiltrated extracts of lignin-degrading peroxidases. Although we observed significant background ABTS oxidation activity in GFP-expressing control extracts, no dimer oxidation was observed in the case of GFP-expressing extracts or in the absence of a peroxide generating enzyme, indicating that dimer conversion was specific to the combined activity of heterologously-produced lignin-degrading enzymes and not other proteins native to the *N. benthamiana* apoplast extract (**Figs. S8 and S9**). Accumulation of guaiacol as the second product resulting from dimer cleavage was not observed, likely due to its subsequent polymerization by the peroxidase. Total product formation occurred to a greater extent under conditions favoring substrate oxidation directly at the enzyme surface^30^ as compared to via Mn(III) as a diffusible mediator^31^ (see PE-*vpl2* & CS-*lip1* at pH 3.5 & 4.0 without Mn(II), vs. PC-*mnp1* & PE-*vpl2* at pH 4.5 with the inclusion of Mn(II) and malonic acid as a chelator, respectively) (**Figs. 3a and S10**). Of the different observed products, C-C bond cleavage was also more limited in the case of MnP-like enzymes (*e.g.* PC-*mnp1*), as indicated by the proportion of veratraldehyde detected relative to dehydrodimer (**Fig. S10**). In comparing the activity of peroxidases without added Mn(II), products accumulated to a similar extent regardless of the enzyme used to generate peroxide *in situ* (**Fig. 3a**, PE-*vpl2* & CS-*lip1* with AN-*gox*, TV-*pox*, or PE-*aao*).

**Figure 3.**
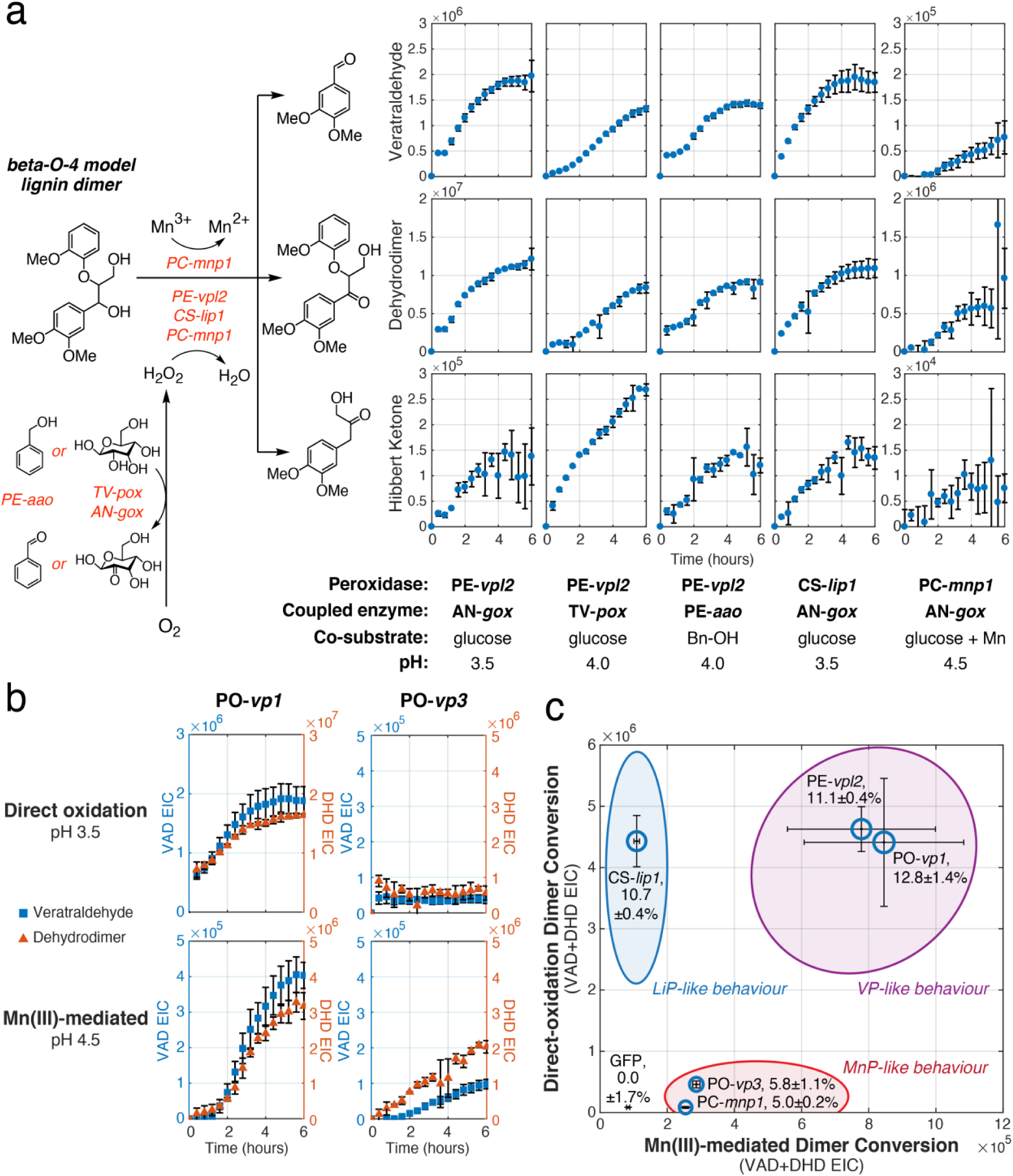
Cleavage of a model β-O-4 lignin dimer using heterologous ligninases produced by *N. benthamiana.* **a)** Diafiltrated extracts of a versatile peroxidase (PE-*vpl2*), a lignin peroxidase (CS-*lip1*) or a manganese peroxidase (PC-*mnp1*) were coupled with peroxide-generating sugar oxidases (TV-*pox* or AN-*gox*) or aryl alcohol oxidase (PE-*aao*(FX9)) in the presence of corresponding co-substrates and under the indicated pH conditions. PE-*vpl2* was found to have negligible activity toward benzyl alcohol (Bn-OH) but was readily accepted by PE-*aao* as a substrate, enabling this coupling reaction to proceed without interference between the two enzymes (**Fig. S13**). Decomposition products (veratraldehyde, dehydrodimer, and Hibbert ketone) were tracked over the course of the reaction using LC-MS. **b)** Dimer oxidation by GOx-coupled diafiltrated extracts of two isoforms of versatile peroxidase from *P. ostreatus* (PO-*vp1* and PO-*vp3*) were compared to PE-*vpl2*, CS-*lip1*, and PC-*mnp1* under direct oxidation and Mn oxidation conditions. **c)** Activity phase diagram of white-rot lignin-degrading peroxidases. The extent of dimer conversion to veratraldehyde (VAD) and dehydrodimer (DHD) in reactions containing peroxidase isozymes coupled to glucose oxidase was measured under direct-oxidation conditions (pH 3.5) and Mn(III)-mediated conditions (pH 4.5 with 1 mM MnSO_4_). The extent of dimer cleavage was calculated as the proportion of veratraldehyde (VAD) EIC relative to the sum of veratraldehyde (VAD) EIC and dehydrodimer (DHD) EIC. The maximum dimer cleavage extent under either condition is displayed in the peroxidase isozyme labels and represented by marker size. Data points represent the average of three independent reaction replicates with error bars calculated as one standard deviation.

Our collection of enzymes successfully produced from *N. benthamiana* included two isoforms of versatile peroxidase from *P. ostreatus*, PO-*vp1* and PO-*vp3*. This provided an opportunity to investigate the specialized roles of the numerous isozymes involved in fungal lignin degradation by analyzing their capacity for model dimer oxidation under different conditions. We observed that coupled reactions involving PO-*vp1* produced substantially greater conversion of lignin dimer compared to PO-*vp3*, especially under direct oxidation conditions (**Fig. 3b**). Comparing the maximal dimer cleavage extent across oxidation conditions, we found that C-C bond cleavage by PO-*vp1* also exceeded that of PO-*vp3* (**Figs. 3c and S10**). Together, these findings suggest that PO-*vp1* is better tailored for oxidation as well as cleavage of this type of lignin bond compared to its relative PO-*vp3* in any oxidation mode.

We expanded this approach to all isozymes tested in the coupled context. Our technique involving high-throughput LC-MS reaction analysis paired with rapid soluble production of multiple isozymes through the *N. benthamiana* platform enabled us to construct a quantitative activity phase diagram portraying the characteristic behavior of isozymes on the model lignin dimer substrate (**Fig. 3c**). We constructed this diagram using dimer conversion extents of each of the isozymes under direct and Mn(III)-mediated oxidation conditions, and identified three behavioral regimes of dimer oxidation, consistent with the three identified families of white-rot lignin-degrading peroxidases. However, unlike previous categorization of isozymes based on activity towards substrates that do not model lignin linkages, such as veratryl alcohol and Mn(II),^9^ this diagram reflects their specific behavior on a major lignin bond type, suggesting that isozymes such as PO-*vp3* exhibit MnP-like interactions with lignin despite its classification as a versatile peroxidase.

### Coupling condition optimization

In an attempt to inform engineering efforts aimed at lignin deconstruction, we sought to identify reaction conditions that maximized conversion of lignin dimer. Given the susceptibility of lignin-degrading peroxidases to inactivation by excess peroxide, we reasoned that optimized coupling of peroxide-generating enzymes would yield greater dimer conversion over the course of the assay compared to exogenously added peroxide in an uncoupled system. Our results indicated that model dimer conversion occurred most effectively under conditions favoring direct oxidation (**Fig. 3c**), irrespective of the peroxide-generating enzyme used (**Fig. 3a**). Accordingly, we selected the coupling of FPLC-purified PE-*vpl2* and commercially-purified AN-*gox* (Sigma) for optimization and focused on the peroxide generation rate as a function of AN-*gox* concentration.

For a given amount of PE-*vpl2*, coupling to glucose oxidase was a more effective strategy for achieving dimer conversion than direct addition of peroxide, enabling approximately four-fold greater product accumulation (**Fig. 4ab**). In the coupled enzyme reaction, the total product formation seems to be limited only by the stability of glucose oxidase under the conditions of the assay (**Fig. S11**). Coupling successfully overcame the limit on conversion in the case of direct peroxide addition, where increasing peroxide concentration did not result in additional product formation, likely due to peroxidase inactivation (**Fig. 4ab, bottom**). We identified an optimum glucose oxidase concentration, below which the rate of the coupled reaction limited dimer conversion within the lifetime of glucose oxidase, and beyond which peroxide likely was generated in excess and resulted in peroxidase inactivation as evidenced by rapid loss of activity (**Fig. 4ab, top**). This maximum concentration could be extended through addition of catalase (*data not shown*) as well as increasing the concentration of peroxidase to accelerate the rate of reaction and therefore enable greater substrate conversion within the lifetime of glucose oxidase (**Fig. S11**). Given the importance of maximizing lignin bond breakage in the process of lignin degradation, we also compared the extent of dimer C-C bond cleavage in the two methods of peroxide addition (**Fig. 4d**). We found that at the optimal glucose oxidase concentration, coupling significantly enhanced bond scission compared to the direct addition of peroxide at any concentration, indicating that our coupling approach would be desirable in engineering synthetic routes to lignin deconstruction.

**Figure 4.**
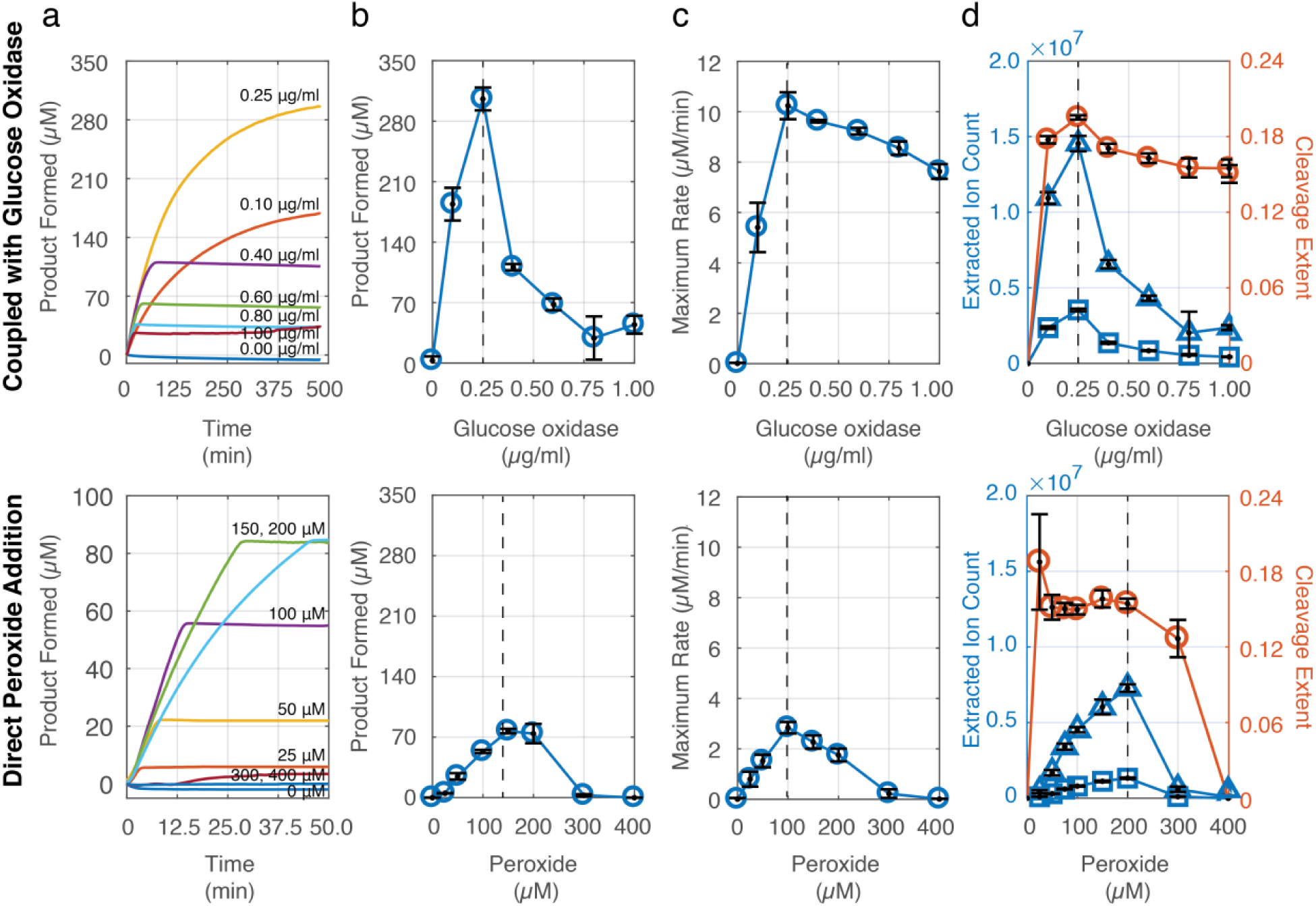
**a)** Kinetic traces of dimer oxidation as a function of glucose oxidase concentration (top) and exogenously added peroxide (bottom). Absorbance was measured at 310 nm and converted to an estimated micromolar aldehyde produced using the molar extinction coefficient for veratraldehyde (9300 1/M 1/cm). Kinetic reactions were performed in triplicate, with a representative time trace shown here. **b)** Maximal dimer conversion achieved over the course of the assay as a function of glucose oxidase concentration (top) or peroxide (bottom), using the molar extinction coefficient for veratraldehyde. **c)** Maximum oxidation rate observed during dimer oxidation as a function of glucose oxidase concentration (top) or peroxide (bottom). **d)** LC-MS analysis of the reaction products produced by dimer oxidation as a function of glucose oxidase concentration (top) or peroxide (bottom). The extracted ion counts (EIC), normalized to the conditions lacking glucose oxidase or peroxide, are shown in blue (*triangles*, dehydrodimer; *squares*, veratraldehyde). The cleavage extent (red) was calculated as the ratio of veratraldehyde EIC to the sum of dehydrodimer and veratraldehyde EIC. All reactions were performed at 25°C and pH 4.0 containing 0.2 μM versatile peroxidase *vpl2* from *P. eryngii* produced and FPLC-purified from apoplast extracts of transgenic *N. benthamiana*. Data points represent the average of three independent reaction replicates with error bars calculated as one standard deviation.

## Discussion

Through the use of *Agrobacterium-*mediated transient expression in *N. benthamiana*, we achieved soluble heterologous production of the majority of basidiomycete enzymes tested (23 of 35). This represents a significant step in overcoming a major obstacle in the study of this class of enzymes and enables rapid combinatorial characterization of their concerted roles in lignin biodegradation. Coupled with LC-MS kinetic analysis, our approach afforded new and technologically valuable insights into the activity of these enzymes in the context of the most commonly occurring linkage in lignin. Whereas previously limitations in the production of lignin-degrading enzymes meant that they were characterized individually, in this study we were able to quantitatively compare the capacity of a set of lignin-degrading peroxidase isoforms to oxidize and cleave this ubiquitous lignin bond under different activity modes. We found that coupling to a peroxide-generating enzyme such as a sugar oxidase enabled greater peroxidase-catalyzed dimer oxidation as well as bond cleavage compared to direct addition of peroxide, suggesting that coupling would be desirable in the context of enzymatic valorization of lignin, especially with peroxide-generating enzymes optimized for greater stability under assay conditions. In this coupled context, we discovered that certain isoforms within a peroxidase family are significantly more proficient in the oxidation and cleavage of this model lignin dimer than others. In the context of natural lignin biodegradation, this suggests some isoforms are more specialized for the initial stages of lignin oxidation, while the activity of other isoforms towards simpler substrates such as Mn and dimethoxyphenol^22^ indicates they may be more involved in downstream oxidation of lignin degradation products. The coupled kinetic characterization approach described here could be readily applied for the study of isozyme specialization towards other commonly-occurring lignin linkages (such as 5-5’ and β-5) through suitable small-molecule model dimers. Combined with our findings described here regarding the β-O-4 dimer, this information about isozyme specialization could serve as initial guidelines for engineering a biochemical route to lignin deconstruction.

The substantially increased capacity of *N. benthamiana* relative to *S. cerevisiae* for the heterologous production of a variety of fungal lignin-degrading enzymes highlights the challenges of selecting an appropriate chassis for these enzymes, especially the heme peroxidases. As synthetic biology for the study of basidiomycete genetics remains in its infancy, the cellular components necessary for successful folding and processing of lignin-degrading enzymes remain unknown. The results shown here in *S. cerevisiae* as well as previous studies in filamentous fungi^10^ suggest that these simpler, genetically tractable organisms lack the specialized cellular environments required for functional expression of lignin-degrading enzymes. For example, we observed hyperglycosylation of lignin-degrading peroxidases in *S. cerevisiae* (**Fig. S12**) compared to single defined glycoforms of *N. benthamiana* apoplast extracts (**Fig. S8**), indicating misrecognition of the nascent protein in the ill-equipped heterologous yeast organism.

The extraordinary oxidative potential of the heme peroxidases in particular also raises questions regarding possible specialized mechanisms of basidiomycete fungi and presumably plant species to handle the oxidative stress resulting from the production of these peroxidases. In particular, the relative ease of heterologous production of different manganese peroxidases compared to lignin peroxidases in *N. benthamiana* (**Fig. 2d**) suggests protein-level determinants of successful folding and export. The development of *N. benthamiana* as a platform amenable to genetic engineering and proficient at lignin-degrading enzyme production affords the opportunity to develop hypotheses aimed at uncovering key cellular components for each class of enzyme, which could inform tailoring of other heterologous production platforms for large-scale production of these biotechnologically important enzymes.

The expression of basidiomycete lignin-degrading enzymes in a lignified plant host also enables *in planta* studies focused on biomass valorization. The transient expression platform described here could be harnessed to trigger lignin decomposition at late stages of plant growth, for instance to facilitate improved biosaccharification by increasing enzyme accessibility of cellulose, augmenting previous work involving bacterial lignin-degrading enzymes^38,39^. This inducible strategy is advantageous in that plant fitness effects would be mitigated relative to existing strategies^40^ aimed at reducing and/or modifying lignin biosynthesis. Through metabolomic analysis and lignin visualization techniques, our platform also provides a framework for characterization of heterologous lignin-modifying enzymes in the context of the plant host itself. Analogous to our *in vitro* approach described here, rapid combinatorial expression of these enzymes *in planta* would provide accelerated insight into their roles within the more complex and therefore more informative environment. This expanded understanding, combined with an ever-evolving array of synthetic biology tools for plant hosts, would substantially make more accessible biorefineries aimed at producing valuable products from the millions of tons of lignocellulose available globally each year.

## Methods

### Chemicals and reagents

Veratryl alcohol (3,4-dimethoxybenzyl alcohol, A13396) and ABTS (2,2’-azino-bis[3-ethylbenzothiazoline-6-sulfonic acid] diammonium salt, J65535) were purchased from Alfa Aesar (Haverhill, MA, USA). β-O-4 dimer (1-[3,4,-dimethoxyphenyl]-2-[2-methoxyphenoxy]propane-1,3-diol, AK-40175) was purchased from Ark Pharm (Arlington Heights, IL, USA). DCIP (dichloroindophenol, D1878) was purchased from Sigma-Aldrich (St. Louis, MO, USA).

Glucose oxidase from *Aspergillus niger* (G2133) and lignin peroxidase from *Phanerochaete chrysosporium* (42603) were purchased from Sigma-Aldrich (St. Louis, MO, USA). Peroxidase from horseradish (31941) was obtained from SERVA (Heidelberg, Germany).

### Protein expression in S. cerevisiae

*S. cerevisiae* strain JHY693^24^ (gift of J. Horecka and A. Chu, Stanford Genomic Technology Center) was used as the background strain for all yeast protein expression. Ligninase genes were codon-optimized (Gen9) for expression in *S. cerevisiae* and synthesized *de novo* using DNA sequences coding for the mature enzymes sourced from UniProt and/or MycoCosm (Joint Genome Institute) databases. For single-copy expression vectors, a pRS415-based cassette (gift of C. Harvey, Stanford Genomic Technology Center) was used; for multi-copy expression vectors, the 2-micron cassette pCHINT2AL^24^ was used (gift of C. Harvey). Inducible expression was driven by the ADH2 promoter of *S. cerevisiae*, and ER targeting and secretion of ligninases was achieved by N-terminal fusion to an evolved variant of the α-mating-factor prepropeptide of *S. cerevisiae*^28^. Yeast transformation was carried out using the Frozen-EZ Yeast Transformation II Kit (Zymo Research). Transformant selection was performed using synthetic defined media plates deficient in leucine. Single colonies were picked into 0.5 ml SD-leu media in a 96-well culture plate and incubated overnight with orbital shaking (425 rpm, 30 C). After centrifugation (600xg, 10 min), the supernatant was removed, and the cell pellets resuspended in supplemented YPEG media (2% ethanol, 3% glycerol, 70 mM potassium phosphate pH 6.0; for heme peroxidase production, 0.01 mM hemin + 1 mM CaCl_2_; for cellobiose dehydrogenase production, 0.01 mM hemin; for laccase production, 2 mM CuSO_4_; no additional supplements for pyranose oxidase or aryl alcohol oxidase production) and incubated for 48 hours with orbital shaking (425 rpm, 20 C). After centrifugation (600xg, 10 min), the culture supernatant was used for subsequent activity assays at 10% v/v.

### Protein expression in N. benthamiana

*Agrobacterium*-mediated transient expression was performed as described before^41^. Genes encoding mature ligninases were cloned from *S. cerevisiae* vectors above into the pEAQ-HT expression cassette^29^. The signal peptide of the dirigent protein of *Sinopodophyllum hexandrum* (UniProt A0A059XIK7, residues 1-27) was used to direct protein export to the apoplast. Expression of ligninases was driven by a 35S promoter from cauliflower mosaic virus (CaMV) and a 5’UTR from cowpea mosaic virus (CPMV). *Agrobacterium* strains harboring the expression cassette and a p19-silencing plasmid were grown on dual-selecting kanamycin-gentamycin LB plates (30 C, 2 days), incubated in induction buffer (150 μM acetosyringone, 10 mM MgCl_2_, 10 mM sodium succinate, pH 5.6; 4-5 hours) before being infiltrated into the three youngest leaves of 5- to 7-week-old *N. benthamiana* plants at an OD_600_ of 0.3 in induction buffer. 4 days post-infiltration, apoplastic contents were extracted as previously described with modifications^27^. Leaves were harvested and submerged in ice-cold extraction buffer (0.1 M sodium acetate, 0.3 M NaCl, pH 4.5); it was observed that 2-(N-morpholino)ethanesulfonic acid (MES) buffer has an inhibitory effect on peroxidase activity (**Fig. S14**). Leaves were subjected to vacuum cycles (>650 mmHg, 3x 5 min) to infiltrate the leaves with buffer. Leaves were individually placed on a piece of Parafilm and rolled around a pipette tip. This assembly was inserted into a 5-ml plunger-less syringe contained in a 15-ml conical centrifuge tube and centrifuged (1600xg, 10 min, 4°C) to produce the apoplastic extract, which was further clarified via centrifugation to remove any plant debris (21000xg, 10 min, 4°C). In experiments featuring diafiltrated apoplast extracts, extracts were diafiltrated at least 600-fold with 20 mM sodium tartrate, pH 4.5, 10% *v/v* glycerol through Amicon Ultra-4 10-kDa MWCO centrifugal filters units (EMD Millipore). For sample storage, 1 volume of 40% glycerol was added to 3 volumes extract and stored at -80°C.

### Enzyme activity testing

ABTS activity assays were performed using 4 mM ABTS, 100 μM H_2_O_2_, 50 mM sodium tartrate, pH 3.5. Assays for Mn-dependent ABTS oxidation were performed using 4 mM ABTS, 100 μM H_2_O_2_, 1.0 mM MnSO_4_, and 50 mM sodium malonate, pH 4.5. ABTS oxidation kinetics were observed at 414 nm (extinction coefficient 36000 1/M 1/cm) using a Synergy HTX plate reader at 25°C. Veratryl alcohol activity^30^ was measured as the production of veratraldehyde at 310 nm (extinction coefficient 9300 1/M 1/cm) using 20 mM veratryl alcohol, 100 μM H_2_O_2_, 50 mM sodium tartrate, pH 3.5, at 25°C. Manganese-dependent activity^31^ was measured by Mn(III)-malonate complex formation using 1.0 mM MnSO_4_ and 100 μM H_2_O_2_ in 50 mM sodium malonate (270 nm, 11590 1/M 1/cm) at 25°C. Cellobiose dehydrogenase activity was measured at 522 nm using 10% *w/v* cellobiose, 0.3 mM dichloroindophenol, and 50 mM sodium tartrate, pH 5.0, at 25°C. Pyranose oxidase activity was measured by coupling to ABTS as above with the inclusion of 1 μg commercial horseradish peroxidase (HRP) and 2% *w/v* D-glucose in 50 mM sodium acetate, pH 6.0. For all assays, 1 unit of activity is defined as 1 μmol of observable product per liter per minute, and activities are calculated as the maximum observed rate during the initial phase of the enzyme assays.

### LC-MS kinetic analysis of dimer oxidation

All reactions contained 20 mM β-O-4 dimer and peroxidase-containing diafiltrated extract from *N. benthamiana* to 0.2 µM total heme content as determined by the pyridine hemochromagen assay^42^. Glucose oxidase assays contained 0.4% D-glucose and either 1.0 ng/μl glucose oxidase and 50 mM sodium tartrate pH 3.5, or 0.574 ng/μl glucose oxidase and 50 mM sodium malonate pH 4.5 with 1.0 mM MnSO_4_, where the glucose oxidase concentration was adjusted between the two pH conditions to keep the rate of peroxide generation constant. Aryl alcohol oxidase assays contained 10 mM benzyl alcohol, 40 U/L (HRP-coupled ABTS activity) of diafiltrated extract of PE-*aao*(FX9) from *N. benthamiana*, and 50 mM sodium tartrate pH 4.0. Pyranose oxidase assays contained 0.4% *w/v* D-glucose, 10 U/L (HRP-coupled ABTS activity) of diafiltrated supernatant of TV-*pox* from *S. cerevisiae*, and 50 mM sodium tartrate pH 4.0. Reactions were clarified (21000xg, 5 min) and initiated by the addition of peroxide-generating enzyme.

Model lignin dimer LC-MS kinetic assays were performed using an Agilent 6545 UHPLC Q-TOF running in positive mode with a 6-minute water-acetonitrile gradient (A: water + 0.1% formic acid, B: acetonitrile + 0.1% formic acid: 0 min, 95% A; 0.2 min, 95% A; 3.65 min, 37.5% A; 3.66 min, 5% A; 4.11 min, 5% A; 4.15 min, 95% A; 5.18 min, 95% A; flow rate 0.8 ml/min) on an Agilent RRHD EclipsePlus 95Å C18 column (2.1 x 50 mm, 1.8 µm, 1200 bar). Reaction product profiles were measured every 24 minutes by 1 µl direct injection of reaction vials, which were maintained at 22°C in the autosampler. Extracted ion counts (EIC) were obtained using the ‘Find by Formula’ function in Agilent MassHunter Qualitative Analysis software, using 35 ppm mass tolerance, 35, 500, and 35 ppm symmetric expansion of values for chromatogram extraction for C_9_H_10_O_3_ (veratraldehyde), C_18_H_20_O_6_ (dehydrodimer), and C_11_H_14_O_4_ (Hibbert ketone), respectively. Possible charge carriers and neutral losses were specified as -electron, +H, +Na, +K,+NH_4_, and -H_2_O.

### LC-MS analysis of dimer cleavage extent by peroxidase isozymes

Reactions contained 20 mM β-O-4 dimer and 0.4% *w/v* D-glucose. Reactions assaying direct substrate oxidation contained 50 mM sodium tartrate, pH 3.5, and 1.0 ng/μl glucose oxidase; reactions assaying Mn-mediated substrate oxidation contained 50 mM sodium malonate, pH 4.5, 1 mM MnSO_4_ and 0.574 ng/μl glucose oxidase (adjusted to keep reaction rate similar). The amounts of diafiltrated extracts of PO-*vp1* and PO-*vp3* used in the reactions were normalized to the Mn activity of PC-*mnp1* at a reaction concentration of 0.2 µM (total heme content; ∼6 U/L). The amounts of diafiltrated extracts of PE-*vpl2* and CS-*lip1* used in the reaction were normalized to PC-*mnp1* by total heme content. Diafiltrated extract of GFP-expressing *N. benthamiana* was used as a negative control at 1% *v/v* (total heme content ∼ 0.07 µM) with the addition of 33.3 ng/μl commercial horseradish peroxidase in order to prevent peroxide accumulation. Reactions were clarified (21000xg, 5 min) prior to initiation by addition of glucose oxidase or hydrogen peroxide. After a nine-hour incubation at room temperature, samples were moved to the LC-MS autosampler maintained at 10°C and analyzed by 1 µl direct injection of the reaction contents on an Agilent 6545 Q-TOF running in positive mode with a 6-minute water-acetonitrile gradient (as above) and an Agilent RRHD EclipsePlus 95Å C18 column (2.1 x 50 mm, 1.8 µm, 1200 bar). EIC values were obtained as above.

### Coupled condition optimization

Reactions contained 20 mM β-O-4 dimer, 50 mM sodium tartrate, pH 4.0, and 330 U/L ABTS activity of FPLC-purified^43^ PE-*vpl2* heterologously produced in *N. benthamiana*. Coupled reactions additionally contained 0.4% *w/v* D-glucose. Absorbance corresponding to the formation of dehydrodimer and veratraldehyde was measured at 310 nm using a Synergy HTX plate reader and converted to an estimate of total aldehyde produced using the molar extinction coefficient for veratraldehyde (9300 1/M 1/cm). Reactions were initiated by the addition of peroxide or glucose oxidase.

After completion, 1 μl of the reaction was injected on a 6545 Agilent UHPLC Q-TOF running in positive mode with an 8-minute water-acetonitrile gradient (0 min, 95% A; 0.2 min, 95% A; 5.65 min, 37.5% A; 5.66 min, 5% A; 6.11 min, 5% A; 6.15 min, 95% A; 7.18 min, 95% A; A: water + 0.1% formic acid, B: acetonitrile + 0.1% formic acid; flow rate 0.8 ml/min) on an Agilent RRHD EclipsePlus 95Å C18 column (2.1 x 50 mm, 1.8 µm, 1200 bar). EIC values were obtained as above.

## Acknowledgements

The authors would like to thank the following members of the Stanford Genome Technology Center for yeast vectors and strains, and for invaluable discussions: Joseph Horecka, Angela Chu, Colin Harvey, Bob St. Onge, and Sundari Suresh. The authors thank Russell Jingxian Li, Warren Lau, Amy Calgaro-Kozina, Curt Fischer, Ryan Nett, Kevin Smith, Kai Yuet, and Corinna Brendel for helpful discussions, experimental assistance, and manuscript feedback. The authors thank Steven Hallam for collaboration on a JGI DNA Synthesis User Proposal. Work conducted by the U.S. Department of Energy Joint Genome Institute, a DOE Office of Science User Facility, is supported under Contract No. DE-AC02-05CH11231. This research was supported by U.S Department of Energy Early Career Award Grant No. DE-SC0014112.

## Author Contributions

N.K. and E.S. conceived and designed the experiments. N.K. performed the experiments. Y.Y. and S.D. provided *E. coli* strains harboring DNA used in the experiments. N.K. and E.S. analyzed the data and discussed results. N.K., Y.Y., and E.S. wrote the manuscript.

## Corresponding author

Correspondence to Elizabeth Sattely.

