## Supplemental Information for "A plant host enables the heterologous production and combinatorial study of fungal lignin-degrading enzymes"

### Supporting Information

### Table of Contents


**Figure S1.** BJ5465 versus JHY693 yeast secretion performance
**Figure S2.** Testing of promoters and ER signal peptides in *S. cerevisiae*
**Figure S3.** Laccase and pyranose oxidase production in *S. cerevisiae* and *N. benthamiana*
**Figure S4.** ABTS activity of white-rot peroxidases tested in *S. cerevisiae*
**Figure S5.** Yeast supernatant inhibition of commercial LiP activity on veratryl alcohol
**Figure S6.** Schematic of enzyme extraction from *N. benthamiana*
**Figure S7.** Testing of different ER signal peptides for *PE-vpl2* production in *N. benthamiana*.
**Figure S8.** Total protein gel and Western blotting of enzymes produced in *N. benthamiana*
**Figure S9.** Negative controls for coupling experiments
**Figure S10.** Product formation and dimer cleavage extent by direct and Mn(III)-mediated oxidation
**Figure S11.** Glucose oxidase stability as a function of pH
**Figure S12.** Western blotting of enzymes secreted by *S. cerevisiae*
**Figure S13.** PE-aao(FX9) and PE-vpl2 activities on benzyl alcohol as a substrate
**Figure S14.** Inhibitory effects of MES buffer on veratryl alcohol oxidation by commercial LiP

**Table S1.** Heme concentration of diafiltrated extracts
**Table S2.** List of strains used
**Table S3.** List of vectors used
**Table S4.** List of genes used
**Table S5.** List of ER signal peptides used
**Table S6.** List of antibody epitope tags used

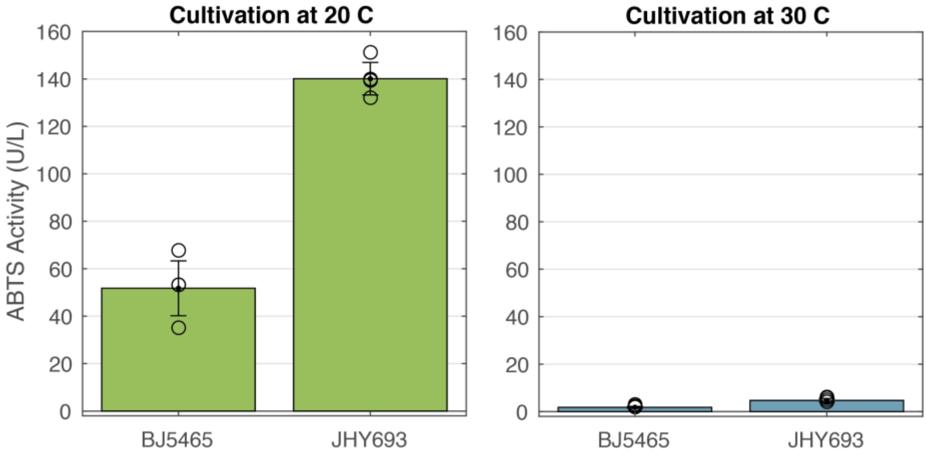


**Figure S1. BJ5465 versus JHY693 yeast secretion performance.** The production of horseradish peroxidase (HRP) was compared in *S. cerevisiae* strains BJ5465[6] and JHY693[2] at 20 and 30 degrees Celsius. ABTS activity was determined as described in Methods.

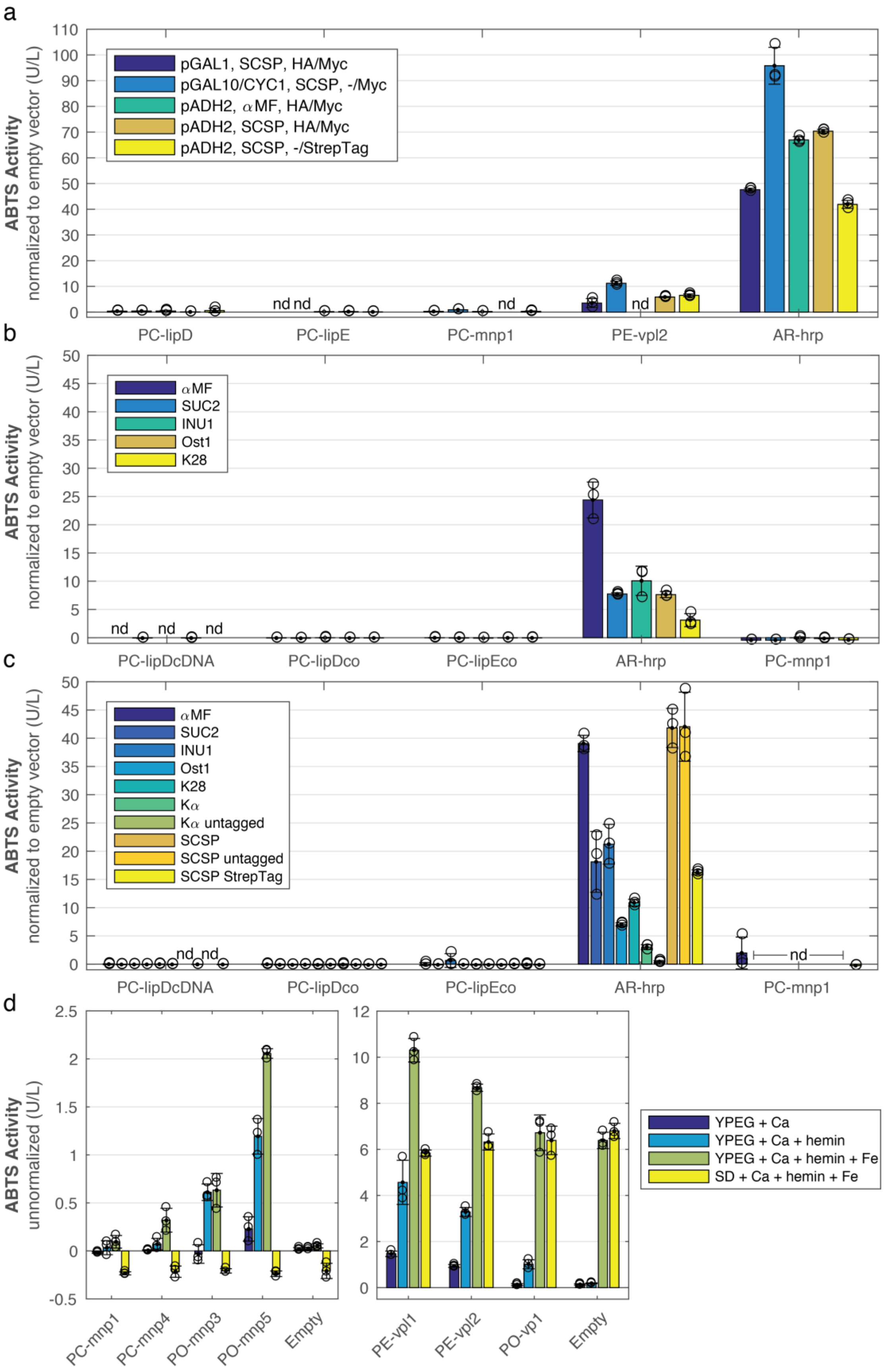

a
ABTS Activity
normalized to empty vector (U/L)
pGAL1, SCSP, HA/Myc
pGAL10/CYC1, SCSP, -/Myc
pADH2, αMF, HA/Myc
pADH2, SCSP, HA/Myc
pADH2, SCSP, -/StrepTag
nd
PC-lipD
PC-lipE
PC-mnp1
PE-vpl2
AR-hrp
b
αMF
SUC2
INU1
Ost1
K28
PC-lipDcDNA
PC-lipDco
PC-lipEco
AR-hrp
PC-mnp1
c
αMF
SUC2
INU1
Ost1
K28
Kα
Kα untagged
SCSP
SCSP untagged
SCSP StrepTag
d
ABTS Activity
unnormalized (U/L)
PC-mnp1
PC-mnp4
PO-mnp3
PO-mnp5
Empty
PE-vpl1
PE-vpl2
PO-vp1
YPEG + Ca
YPEG + Ca + hemin
YPEG + Ca + hemin + Fe
SD + Ca + hemin + Fe

**Figure S2. Testing of promoters and ER signal peptides in *S. cerevisiae.* a)** Comparison of promoter, signal peptides, and antibody affinity tags on production of lignin-degrading peroxidases from *S. cerevisiae* from high-copy 2µ expression cassettes. **b)** Comparison of signal peptide choice in the production of peroxidases in *S. cerevisiae* from the low-copy CEN/ARS expression cassette[1] pRS415-ADH2. **c)** Comparison of signal peptide choice in the production of peroxidases in *S. cerevisiae* from the high-copy 2µ expression cassette pCHINT2AL[2]. **d)** Optimization of media conditions for peroxidase production from pL231 expression cassette in *S. cerevisiae*. YPEG, rich media; SD, synthetic defined media. Signal peptides: αMF, alpha-mating factor, evolved variant appS4[3]; SUC2, *S. cerevisiae* invertase; INU1, *K. marxianus* inulinase; Ost1, *S. cerevisiae* pre-*Ost1*-pro-αMF fusion[4]; K28, K28 killer toxin; Kα, killer toxin alpha subunit; SCSP, synthetic consensus signal peptides[5]. PC, *P. chrysosporium*; PE, *P. eryngii*; PO, *P. ostreatus*; AR, *A. rusticana*; cDNA, sequence from cDNA; co, sequence codon-optimized for expression in *S. cerevisiae*. Antibody affinity tags: HA, hemagglutinin; Myc: c-myc. nd: not determined. ABTS activity was determined as described in Methods. Error bars represent one standard deviation in activity levels of three biological replicates.

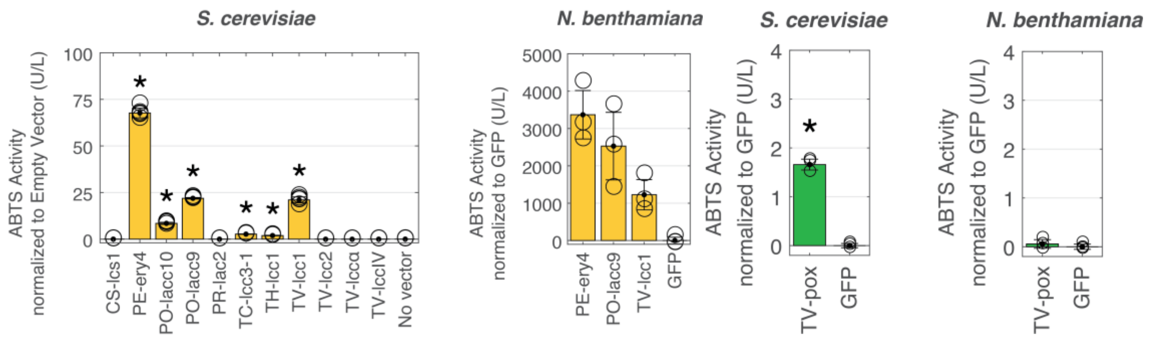


**Figure S3. Laccase and pyranose oxidase production in *S. cerevisiae* and *N. benthamiana*. a)** ABTS activity of supernatant of *S. cerevisiae* and of crude apoplast extracts of *N. benthamiana* plants producing laccases. Error bars for S. cerevisiae measurements represent one standard deviation in activity determined from five biological replicates; asterisks indicate statistical significance relative to a no-vector control ($p < 0.05$). Error bars for N. benthamiana represent one standard deviation in activity determined from three individual leaves as biological replicates. **b)** ABTS activity of supernatant of *S. cerevisiae* and of crude apoplast extracts of *N. benthamiana* plants producing a pyranose oxidase from *T. versicolor* (TV-*pox*). Error bars for represent one standard deviation in activity determined from three biological replicates; asterisk indicates statistical significance relative to a GFP-expressing control ($p < 0.05$).

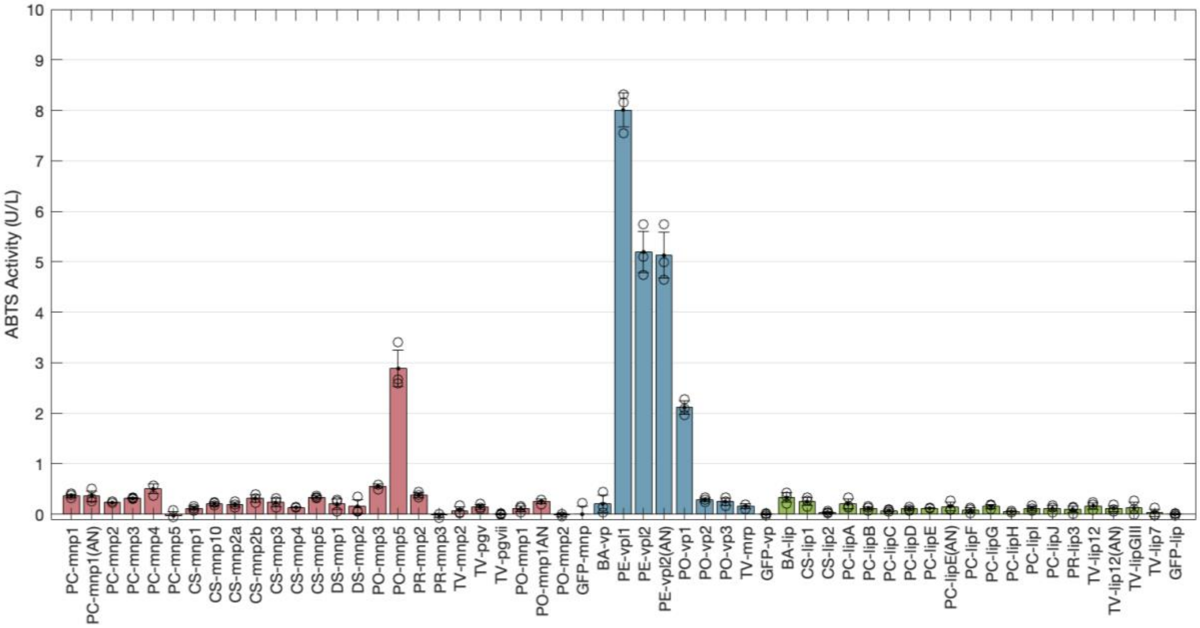


**Figure S4. ABTS activity of all white-rot peroxidases tested in *S. cerevisiae*.** Activity was determined as described in Methods. PC, *P. chrysosporium*; CS, *C. subvermispora*; DS, *D. squalens*; PE, *P. eryngii*; PO, *P. ostreatus*; PR, *P. radiata*; TV, *T. versicolor*; BA, *B. adusta*. GFP corresponds to GFP-expressing strains tested under the corresponding conditions appropriate for the peroxidase type. Error bars represent one standard deviation in activity levels of three biological replicates.

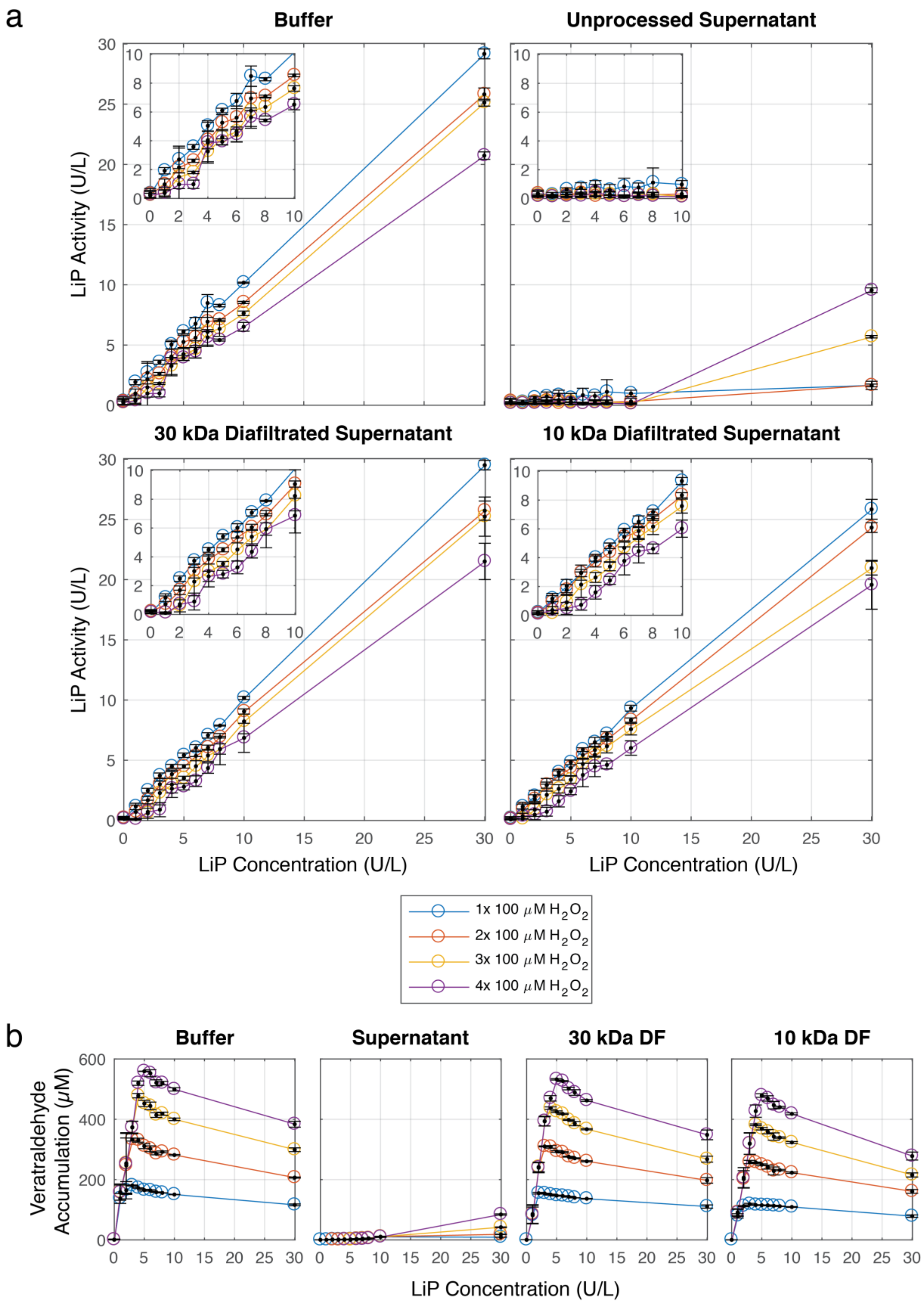

a
Buffer
Unprocessed Supernatant
30 kDa Diafiltrated Supernatant
10 kDa Diafiltrated Supernatant
LiP Activity (U/L)
LiP Concentration (U/L)
1x 100 μM $H_2O_2$
2x 100 μM $H_2O_2$
3x 100 μM $H_2O_2$
4x 100 μM $H_2O_2$
b
Buffer
Supernatant
30 kDa DF
10 kDa DF
Veratraldehyde Accumulation (μM)
LiP Concentration (U/L)

**Figure S5. Yeast supernatant inhibition of commercial LiP activity on veratryl alcohol.** Commercial lignin peroxidase (Sigma) was assayed for activity on veratryl alcohol (VA) in the context of 10%*v/v* buffer, unprocessed supernatant from *S. cerevisiae* expressing GFP, supernatant diafiltrated 10000-fold through 30 kDa and 10 kDa size-exclusion centrifugal filters with 20 mM sodium acetate, pH 6.0. LiP activity on veratryl alcohol was measured by absorbance at 310 nm indicating formation of veratraldehyde as a product ($\varepsilon$ = 9300 1/M 1/cm)[7]. Assays were initiated using 100 µM hydrogen peroxide, which was successively added three more times after full peroxide consumption as indicated by constant absorbance readings. **a)** Observed LiP activity as a function of LiP concentration in the reaction. **b)** Accumulation of veratraldehyde as a function of LiP concentration. Unprocessed yeast supernatant inhibited the formation of veratraldehyde at low LiP concentrations, which was only observed after the third addition of peroxide, presumably after full conversion of inhibiting compounds. Diafiltration eliminated the observed inhibition. Error bars represent one standard deviation of triplicate activity assays.

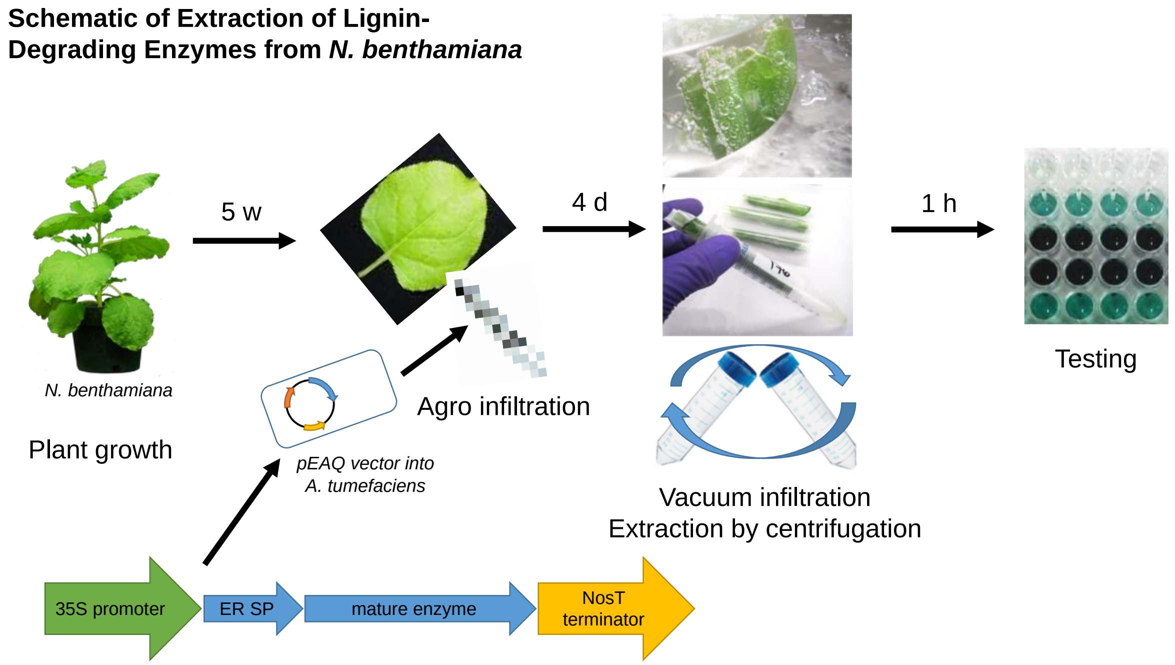


**Figure S6. Schematic of enzyme extraction from *N. benthamiana.*** Crude apoplast extracts are produced from 5-week-old *N. benthamiana* plants transiently transformed with *Agrobacterium tumefaciens* harboring pEAQ expression vectors. Protocol adapted from O'Leary *et al*[8].

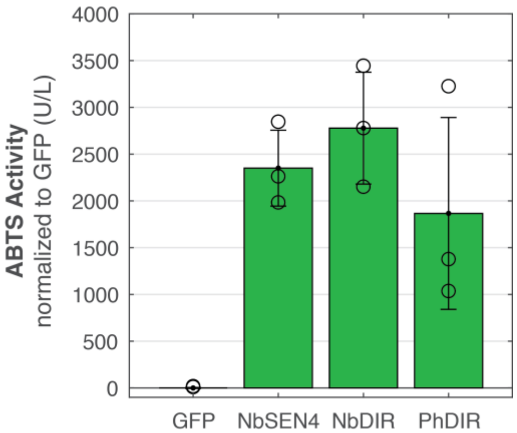


**Figure S7. Testing of different ER signal peptides for *PE-vpl2* production in *N. benthamiana*.** pEAQ expression cassettes harboring the mature *PE-vpl2* sequence were fused with signal peptides derived from xyloglucan endotransglucoslase/hydrolase (*NbSEN4*, UNIPROT A0A1Q1N6K4) of *N. benthamiana* or dirigent protein (*NbDIR*, UNIPROT Q0WYB7) of N. benthamiana, and compared to that of dirigent protein (*PhDIR*) of *P. hexandrum*, which was used for all other expression cassettes in *N. benthamiana*. ABTS activity was measured as described in Methods.

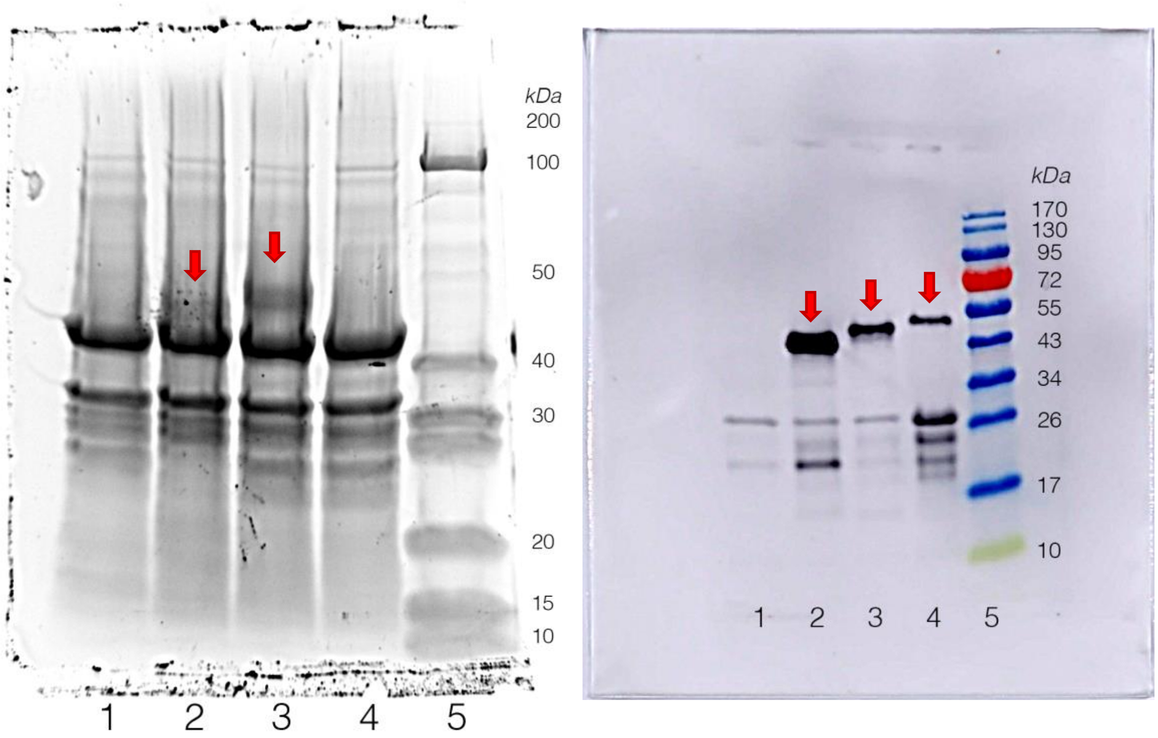


**Figure S8. Total protein gel and Western blotting of enzymes produced in *N. benthamiana*.** *Left:* 5.55 µg of total protein (as measured by Bradford assay[9]) of diafiltrated apoplast extracts of *N. benthamiana* were analyzed by Flamingo (BioRad) staining. Lane 1, GFP control; lane 2, PE-*vpl2*; lane 3, PC-*mnp1*; lane 4, CS-*lip1*; lane 5, protein ladder (Fisher). Red arrows indicate expected bands corresponding to PE-*vpl2* and PC-*mnp1*, respectively; CS-*lip1* presumably too faint to be detected. *Right:* 5 µl of diafiltrated apoplast extracts of *N. benthamiana* were analyzed by Western blotting of C-terminal Myc tag of lignin-degrading peroxidases. Lane 1, GFP control; lane 2, PE-*vpl2*; lane 3, PC-*mnp1*; lane 4, CS-*lip1*; lane 5, protein ladder (Fisher). Red arrows indicate expected bands corresponding to PE-*vpl2*, PC-*mnp1*, and CS-*lip1*. The expected molecular weight of the mature enzymes including affinity tags is between 37, 40 and 38 kDa, respectively.

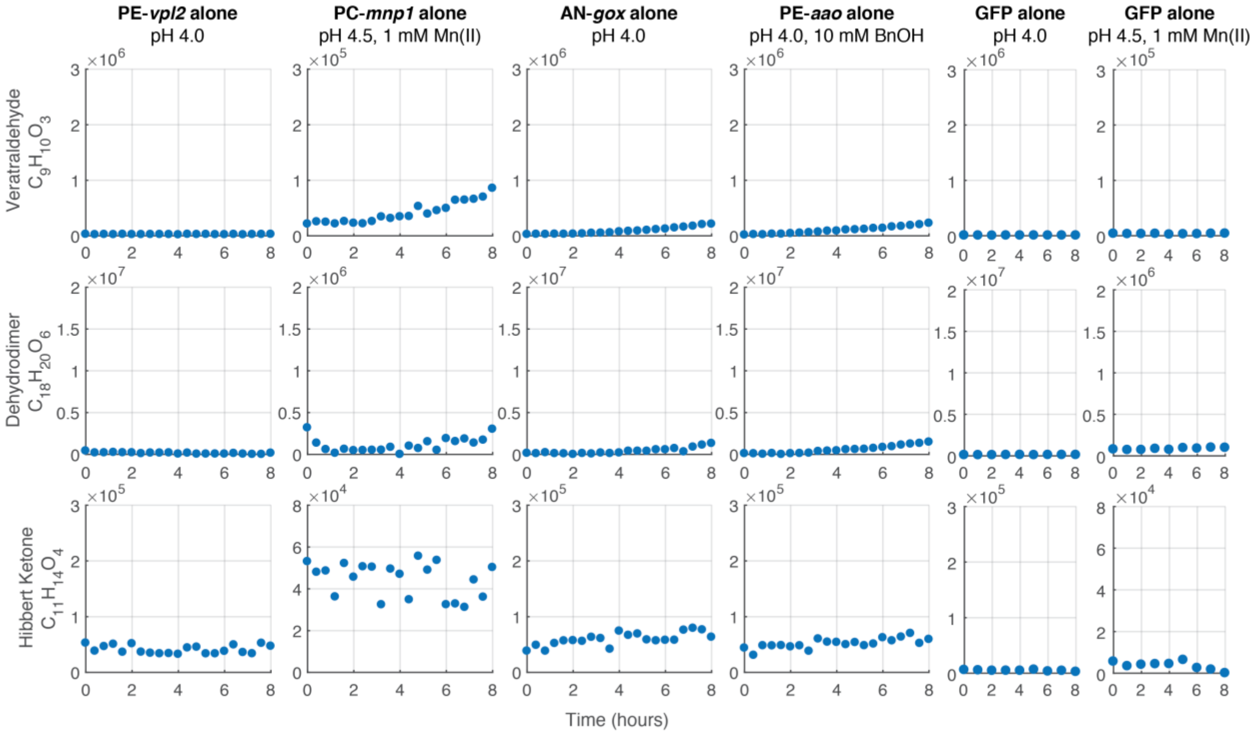


**Figure S9. Negative controls for coupling experiments**. Diafiltrated apoplast extracts of PE-*vpl2* and PC-*mnp1* from *N. benthamiana* were individually assayed for activity towards a model β-O-4 lignin dimer under conditions corresponding to those used in Figure 3 except without the addition of peroxide-generating enzymes. Commercially-available glucose oxidase (AN-*gox*) and diafiltrated apoplast extract of PE-*aao*(FX9) from *N. benthamiana* were tested in the same way without the addition of lignin-degrading peroxidases. Apoplast extract of GFP-expressing *N. benthamiana* was tested in the same way with and without Mn(II), except with lower reaction sampling frequency and 10 mM dimer instead of 20 mM.

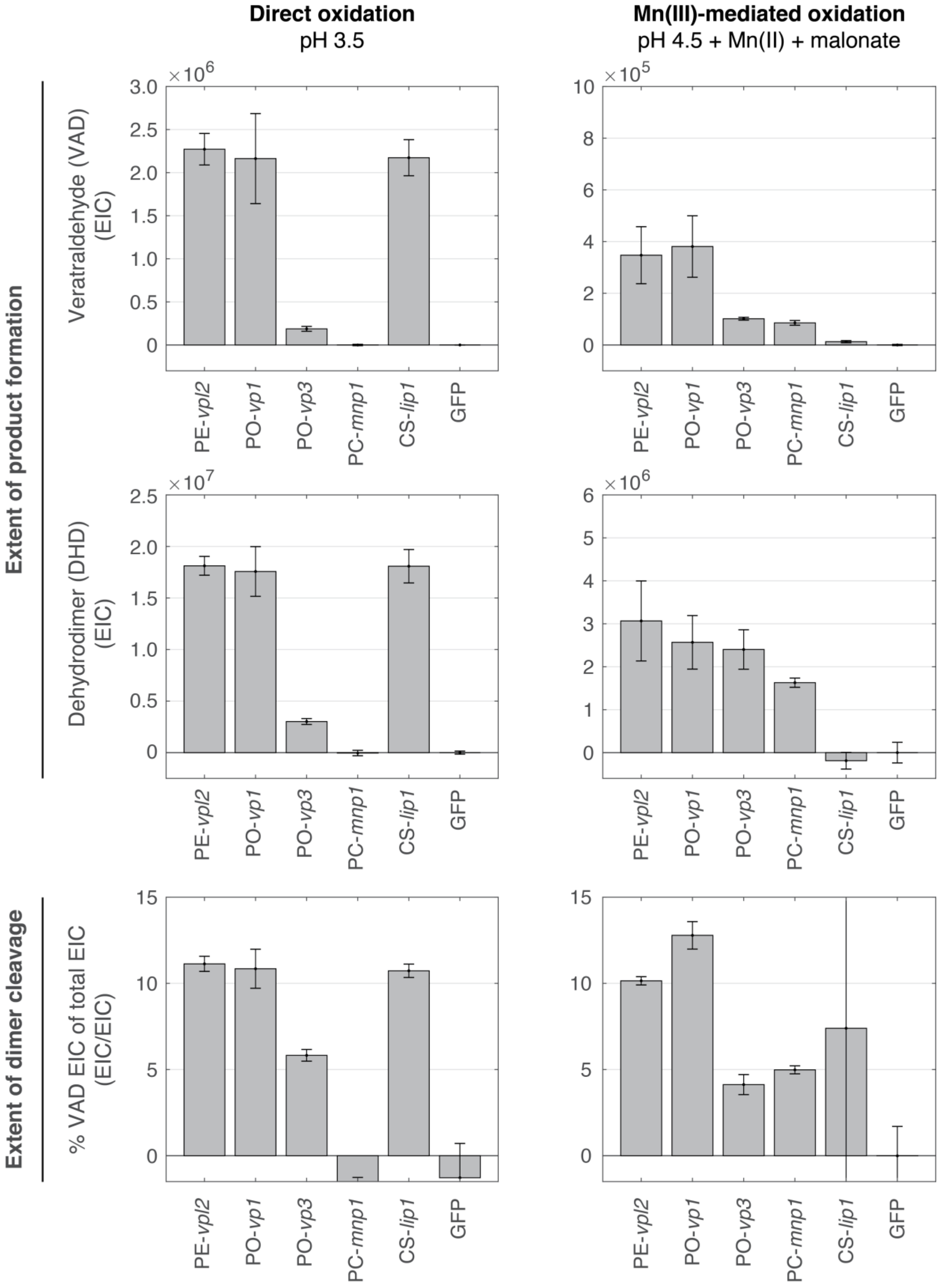

Direct oxidation
pH 3.5
Mn(III)-mediated oxidation
pH 4.5 + Mn(II) + malonate
Extent of product formation
Extent of dimer cleavage
Veratraldehyde (VAD) (EIC)
Dehydrodimer (DHD) (EIC)
% VAD EIC of total EIC (EIC/EIC)
×10^6
×10^5
×10^7
×10^6
PE-vpl2
PO-vp1
PO-vp3
PC-mnp1
CS-lip1
GFP

**Figure S10. Product formation and dimer cleavage extent by direct and Mn(III)-mediated oxidation.** Reactions were performed as described in Methods. Diafiltrated apoplast extracts from heterologous *N. benthamiana* containing peroxidases were coupled with glucose oxidase from *A. niger* in dimer oxidation reactions containing either 50 mM sodium tartrate, pH 3.5, or 50 mM sodium malonate, pH 4.5, and 1 mM $MnSO_4$, representing conditions favoring direct and Mn(III)-mediated oxidation, respectively. Dimer cleavage extent (bottom row) was determined as the proportion of EIC corresponding to veratraldehyde relative to total EIC corresponding to the sum of veratraldehyde and dehydrodimer and is represented as a percentage. The EIC data used for this calculation is net of the EIC detected for the GFP samples, and cleavage extent was calculated for each replicate individually before averaging. Data bars represent the average of three independent replicate reactions, and error bars represent one standard deviation.

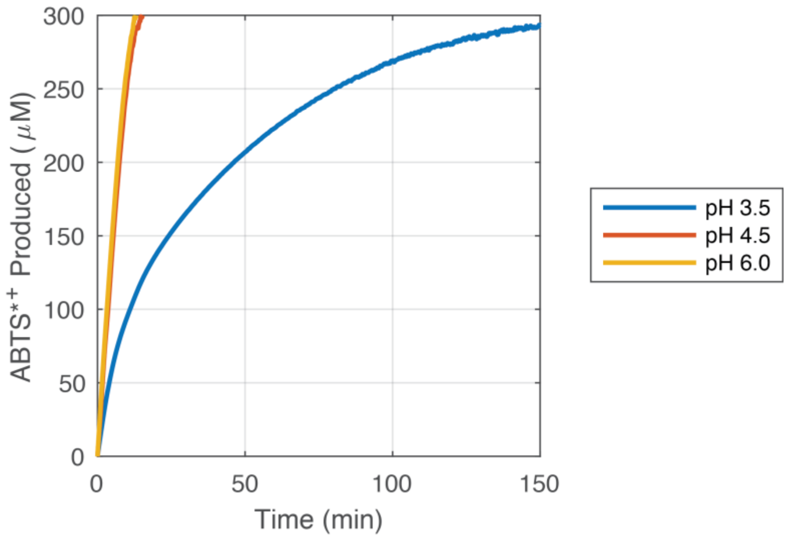


**Figure S11. Glucose oxidase stability as a function of pH.** Glucose oxidase stability was tested as a function of pH with ABTS oxidation as a readout catalyzed by horseradish peroxidase (HRP). Reactions involved 0.574 ng/µl commercial glucose oxidase (Sigma), 4 mM ABTS, 25 ng/µl commercial HRP (Serva), 50 mM sodium tartrate, pH 3.5, or sodium malonate, pH 4.5, or sodium acetate, pH 6.0, and 0.4% *w/v* D-glucose. Reactions were performed at 25 C and ABTS oxidation was measured spectroscopically at 414 nm using an extinction coefficient of 36000 1/M 1/cm. The reactions at pH 4.5 and 6.0 saturated the photodetector while the reaction at pH 3.5 did not, highlighting the instability of glucose oxidase under acidic conditions.

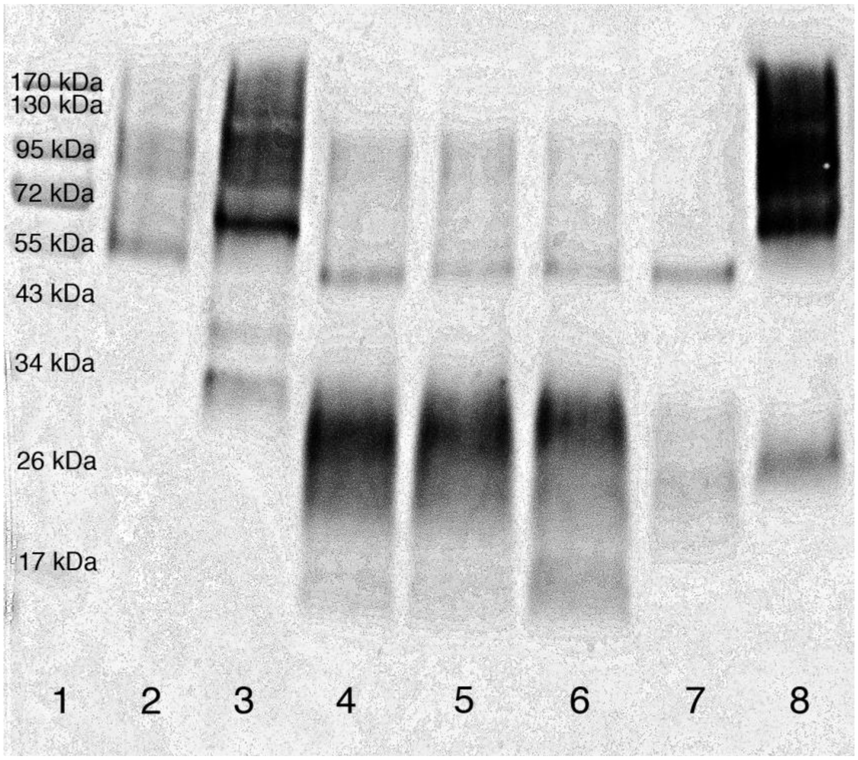


**Figure S12. Western blotting of enzymes secreted by *S. cerevisiae*.** 20 µl of media supernatants of *S. cerevisiae* were analyzed by Western blotting of C-terminal Myc tag of lignin-degrading peroxidases. Lane 1, protein ladder (Fisher); lane 2, BA-*vp*; lane 3, CS-*lip2*; lane 4, PE-*vpl1*; lane 5, PE-*vpl2*; lane 6, PO-*vp1*; lane 7, PO-*vp3*; lane 8, TV-*mrp*. The expected molecular weight of the mature enzymes including affinity tags is between 37 and 38 kDa.

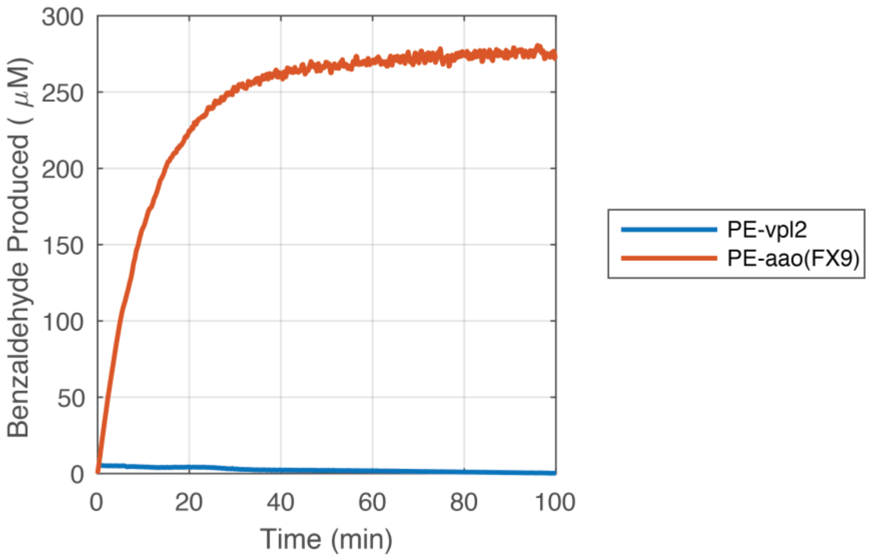


**Figure S13. PE-aao(FX9) and PE-vpl2 activities on benzyl alcohol as a substrate.** Diafiltrated apoplast extracts of *N. benthamiana* expressing PE-*aao*(FX9)[10] or PE-*vpl2* were incubated with 10 mM benzyl alcohol in 50 mM sodium tartrate, pH 3.5 and 100 µM $H_2O_2$ (only for PE-*vpl2*). Benzaldehyde production was measured spectroscopically at 250 nm (ε = 13800 1/M 1/cm)[11]. No benzyl alcohol activity was observed of PE-*vpl2*, whereas benzaldehyde was readily produced by PE-*aao*(FX9). PE-*aao*(FX9) activity decreases over time without full conversion of substrate presumably due to the enzyme's instability at the low pH of the assay.

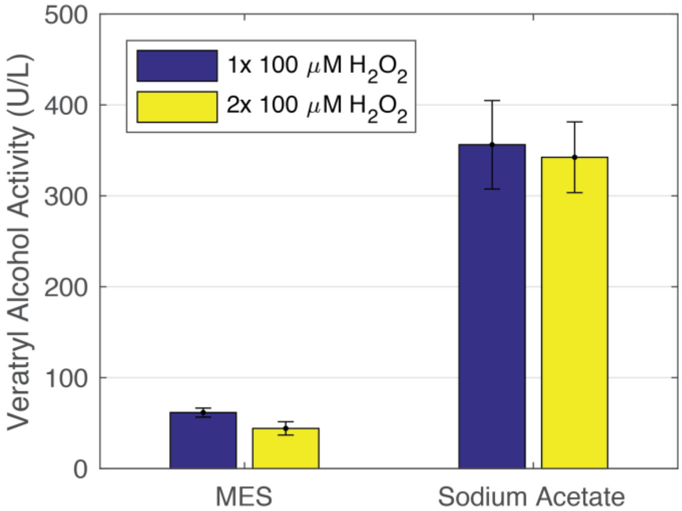


**Figure S14. Inhibitory effects of MES buffer on veratryl alcohol oxidation by commercial LiP.** 0.16 mg/ml commercial lignin peroxidase (Sigma) was tested for activity against veratryl alcohol (2 mM) in the presence of peroxide (0.1 mM) and either 2-(N-morpholino)ethanesulfonic acid (MES) or sodium acetate buffer (13 mM, pH 6.0) using plate reader spectroscopy (see Methods). MES buffer inhibited LiP activity approximately six-fold compared to sodium acetate, and the latter was used for apoplast extraction of enzymes from *N. benthamiana*.

**Table S1. Heme concentration of diafiltrated extracts.** Heme content was measured of diafiltrated apoplast extracts from *N. benthamiana* by the pyridine hemachromagen method[12] using absorbance at 557 nm and a molar extinction coefficient of 34700 1/M 1/cm.

| **PC-*mnp1*** | **PE-*vpl2*** | **CS-*lip1*** | **GFP** |
|---|---|---|---|
| 5.00 µM | 3.93 µM | 4.23 µM | 1.97 µM |

**Table S2. List of strains used.**

| Strain | Species | Genotype | Reference |
|---|---|---|---|
| BJ5465 | *S. cerevisiae* | MATa ura3-52 trp1 leu2-delta1 his3-delta200 pep4::HIS3 prb1-delta1.6R can1 GAL | Ref. 7 |
| JHY693 | *S. cerevisiae* | MATa his3Δ1 leu2Δ0 ura3Δ0 met15Δ0 SAL1+ HAP1+ CAT5(91M) MIP1(661T) MKT1(30G) RME1(INS-308A) TAO3(1493Q) prb1Δ pep4Δ | Ref. 1 |
| GV3103 | *A. tumefaciens* | | |

**Table S3. List of vectors used.**

| Plasmid | Description | Reference |
|---|---|---|
| pRS415-ADH2 | Leu2-$P_{ADH2}$-MCS-$T_{TEF1}$-$ORI_{CEN/ARS}$-AmpR | Dr. Colin Harvey, SGTC |
| pCHINT2AL | Leu2-$P_{ADH2}$-flag-MCS-$T_{TEF1}$-$ORI_{2\mu}$-AmpR | [2] |
| pL131 | Leu2-$P_{ADH2}$-αMF*appS4*-HA-MCS-Myc-$T_{TEF1}$-$ORI_{CEN/ARS}$-AmpR | This study |
| pL231 | Leu2-$P_{ADH2}$-αMF*appS4*-HA-MCS-Myc-$T_{TEF1}$-$ORI_{2\mu}$-AmpR | This study |
| pEAQ | $P_{35S}$-5'$UTR_{CPMV}$-PhDIRSP-MCS-Myc-His6-3'$UTR_{CPMV}$-$T_{NOS}$-KanR | [13] |

**Table S4. List of genes used.**

See attached Excel document.

**Table S5. List of ER signal peptides used.**

| ER Signal Peptide | Protein Sequence | DNA Sequence |
|---|---|---|
| PhDIR | MGGEKAFSFIFLLFVCFFLANLSGSSA | atgggaggagaaaaagctttcagtttcattttcctcctcttcgtgtgcttcttcctagccaacctctctgggtcttcagct |
| NbDIR | MEKLNLILLLSSIAITISSIPFAHA | atggaaaagctaaacctaattctattgctttcctccattgctattaccatatcatcaattccgtttgctcatgcc |
| NbSEN4 | MSCKLVLALMVSAFAIATA | atgtcttgtaaattagtactagctcttatggttagtgcttttgctattgcaactgcg |
| αMFappS4 | MRFPSIFTAVVFAASSALAAPANTTAEDETAQIPAEAVIGYLGLEGDSDVAALPLSDSTNNGSLSTNTTIASIAAKEEGVSLDKREAEA | atgagatttccttcaatttttactgcagttgtattcgcagcatcctccgcattagctgctccagccaacactacagcagaagatgaaacggcacaaattccggctgaagctgtcatcggttacttaggtttagaaggggattccgatgttgctgctttgccattgtccgacagcacaaataacgggtcattgtctacaaatactactattgccagcattgctgctaaagaagaaggggtatctttggataaaagagaggctgaagct |
| SUC2 | MLLQAFLFLLAGFAAKISA | atgcttttgcaagctttccttttccttttggctggttttgcagccaaaatatctgca |
| INU1 | MKLAYSLLLPLAGVSASVINYKR | atgaagttagcatactccctcttgcttccattggcaggagtcagtgcttcagttatcaattacaagaga |

| | | |
|---|---|---|
| pre-Ost1-pro-αMF | MRQVWFSWIVGLFLCFFNVSSAAPVNTTTEDETAQIPAEAVIGYLDLEGDFDVAVLPFSNSTNNGLLFINTTIASIAAKEEGVSLDKREAEA | atgaggcaggtttggttctcttggattgtgggattgttcctatgtttttcaacgtgtcttctgctgctccagtcaacactacaacagaagatgaaacggcacaaattccggctgaagctgtcatcggttacttagatttagaaggggatttcgatgttgctgttttgccattttccaacagcacaaataacgggttattgtttataaatactactattgccagcattgctgctaaagaagaaggggtatctttggataaaagagaggctgaagct |
| K28 | MESVSSLFNIFSTIMVNYKSLVLALLSVSNLKYARG | atggaatccgtcagttccttgttcaacattttctccaccatcatggtcaactacaagtctttggttttggccttgttgtccgtttctaatttgaaatacgctagaggt |
| Kα | MNIFYIFLFLLSFVQGLEHTHRRGSLVKR | atgaatatattttacatatttttgtttttgctgtcattcgttcaaggtttggagcatactcatcgaagaggctccttagtcaaaagg |
| SCSP | MKVLIVLLAIFAALPLALAQPVISTTVGSAAEGSLDKREA | atgaaggttttgattgtcttgttggctatcttcgctgctttgccattggccttagctcaaccggttatttctactaccgtcggttccgctgcagaaggctctttggacaagagagaagct |

**Table S6. List of antibody epitope tags used.**

| Tag | Protein Sequence | DNA Sequence |
|---|---|---|
| Human influenza hemagglutinin (HA) | YPYDVPDYA | tacccatacgacgttccagactacgct |
| c-Myc (Myc) | EQKLISEEDL | gaacaaaagcttatttctgaagaggacttg |
| StrepTag | WSHPQFEK | tggtctcatccacaatttgaaaaa |

| | | |
|---|---|---|
| Myc-His6 | ASEQKLISEEDLNSAVD HHHHHH | gctagcgaacaaaaactcatctcagaagaggatctgaatagcgccgtc gaccatcatcatcatcatcat |
